# Tyk2-mediated signaling promotes the development of autoreactive CD8^+^ CTLs and autoimmune type 1 diabetes

**DOI:** 10.1101/2023.07.14.548984

**Authors:** Keiichiro Mine, Seiho Nagafuchi, Satoru Akazawa, Norio Abiru, Hitoe Mori, Hironori Kurisaki, Yasunobu Yoshikai, Hirokazu Takahashi, Keizo Anzai

**Author notes:** Corresponding author: Keiichiro Mine. Mailing address: Division of Metabolism and Endocrinology, Department of Internal Medicine, Faculty of Medicine, Saga University, 5-1-1, Nabeshima, Saga City, Saga, 849-0937, Japan.

## Abstract

Tyrosine kinase 2 (TYK2), a member of the JAK family, might be a susceptibility gene for type 1 diabetes (T1D), whereas its precise role in autoimmune T1D remains unknown. We showed *Tyk2* deficiency and inhibition suppressed autoimmune T1D development in non-obese diabetic (NOD) mice. Defective IL-12 signaling due to *Tyk2* deficiency in islet-autoreactive CD8^+^ CTLs during their priming reduced T-bet expression, leading to impaired Cxcr3 expression and effector functions against β-cells. *Tyk2* deficient CD8^+^ resident dendritic cells (rDC) exhibited reduced MHC I expression and impaired cross-priming of CTLs. In β-cells, increased expressions of *Fas*, MHC I, and chemokines with age were attenuated by *Tyk2* deficiency. We demonstrated that treatment with BMS-986165, a Tyk2 inhibitor, inhibited the development of CTLs and inflammation in β-cells *in vitro*. BMS-986165 reduced the incidence of diabetes in NOD mice. Thus, we demonstrated that Tyk2-mediated signaling has a critical role in the development of autoreactive CD8^+^ CTLs, inflammation in β-cells, and the pathogenesis of autoimmune T1D.

**Summary:** We demonstrated that Tyk2-mediated signaling plays a critical role in the development of autoreactive CD8^+^ CTLs, inflammation in β-cells, and the pathogenesis of autoimmune T1D. These findings will lead to the development of safety and effective prevention strategies for T1D.

## INTRODUCTION

The immune system is strictly regulated to maintain homeostasis by balancing immune tolerance and immunogenicity, and therefore, a break in this balance leads to autoimmune disorders or increased susceptibility to infectious agents (Horwitz et al., 2019). It is thought that a combination of genetic and environmental factors contribute to the pathogenesis of autoimmune diseases. Multifaceted approaches, although challenging, are needed to uncover the pathogenic mechanisms of autoimmune disease and to develop effective preventive/therapeutic measures against multifactorial diseases (Ellis et al., 2014; Gorman et al., 2017).

T1D is caused by the destruction of insulin-producing pancreatic β-cells and requires lifelong insulin therapy. The number of patients with T1D worldwide in 2021 was approximately 8.4 million and this is expected to increase rapidly because of changes in environment and lifestyle (Gregory et al., 2022; Knip and Simell, 2012). T1D is a multifactorial disease related to genetic and environmental factors with a heterogeneous and complex pathogenesis (DiMeglio et al., 2018). Genetic studies of T1D revealed risk loci associated with immune responses (Onengut-Gumuscu et al., 2015; Chiou et al., 2021; Bradfield et al., 2011). A candidate susceptibility gene of T1D is tyrosine kinase 2 (TYK2) (Onengut-Gumuscu et al., 2015; Bradfield et al., 2011), a member of the Janus kinase (Jak) family. TYK2 is expressed ubiquitously and involves in the signal transduction of type 1 IFNs, IL-23, IL-12, IL-10, and IL-6 (Leitner et al., 2017). When these cytokines bind to cell surface receptors, TYK2 is activated, phosphorylates the intracellular tail of cytokine receptors, and recruits signal transducers and activators of transcriptions (Stats) in concert with other Jaks. The recruited Stats are phosphorylated, dimerized, and translocated to the nucleus to regulate cytokine-specific gene expression and cytokine secretion (O’Shea et al., 2015). These Tyk2-mediated responses confer host defense against microorganisms (Li et al., 2007; Hashiguchi et al., 2014; Izumi et al., 2015). Of note, TYK2-deficient patients had impaired type 1 IFNs and IL-12 signal transduction, resulting in mycobacterial and viral infections (Kreins et al., 2015).

Viruses were reported to be potential environmental factors related to T1D (Mine et al., 2020b; Coppieters et al., 2012; Fukui et al., 2023). Viral infection may contribute to the development of T1D by several mechanisms including direct β-cell destruction, triggering autoimmunity to β-cells, molecular mimicry, and induction of β-cell dedifferentiation (Girdhar et al., 2022; Mine et al., 2020b; Coppieters et al., 2012). We previously reported that reduced *Tyk2* expression in β-cells led to impaired anti-viral defense, increasing the risk of β-cell-tropic virus-induced diabetes in mice (Izumi et al., 2015). Furthermore, we found that a polymorphism in the promoter region of the *TYK2* gene (ClinVar, 440728), which decreased promoter activity, was a risk factor for T1D, particularly in patients with T1D associated with flu-like syndrome and who were autoantibody negative at the onset of disease (Nagafuchi et al., 2015; Mine et al., 2017). Although the role of Tyk2 in virus-induced β-cell destruction is controversial, these observations suggest that Tyk2 might be a host susceptibility gene for virus-induced diabetes.

However, Tyk2-mediated signaling is associated with the pathogenesis of autoimmune diseases. Indeed, the loss-of-function mutations and inhibition of Tyk2 suppressed autoimmune processes in mice and humans (Ishizaki et al., 2014; Burke et al., 2019; Gracey et al., 2020; Yuan et al., 2023). The role of Tyk2 in psoriasis, which predominantly manifests as skin lesions (Greb et al., 2016), is well characterized. In a mouse model of imiquimod-induced psoriasis-like dermatitis, IL-17 and IL-22 secreted from γδ T cells and Th17 upon IL-23 stimulation were involved in skin inflammation and keratinocyte activation, which were improved by Tyk2 inhibitor treatment (Ishizaki et al., 2014; Works et al., 2014; Ishizaki et al., 2011). In patients with psoriasis, oral treatment with a selective Tyk2 inhibitor, BMS-986165 (deucravacitinib), was efficacious (Strober et al., 2023). Furthermore, a Tyk2 inhibitor ameliorated disease progression in murine models of spondyloarthritis, lupus nephritis, and inflammatory bowel disease (Burke et al., 2019; Gracey et al., 2020). These studies demonstrate the involvement of Tyk2 in the development of autoimmune disease and its therapeutic potential, although the precise role of Tyk2 in autoimmune T1D remains unknown.

In this study, we generated *Tyk2* KO non-obese diabetic (NOD) mice, a well-established model of autoimmune T1D (Reed and Herold, 2015), and revealed the mechanisms involved in the Tyk2-mediated pathogenesis of autoimmune T1D. Additionally, we showed the potential efficacy of a Tyk2 inhibitor to prevent the development of autoimmune T1D. These findings demonstrate the role of Tyk2 in the development of autoimmune T1D and provide information that might aid in developing safe and effective prevention strategies for T1D.

## RESULTS

### *Tyk2* deficiency suppresses autoimmune T1D in NOD mice

To determine the role of Tyk2 in autoimmune T1D, we generated *Tyk2* gene KO NOD mice by backcrossing Tyk2KO.B6 mice onto the NOD background for 10 generations. We confirmed all insulin-dependent diabetes susceptibility loci were of NOD origin (Figure S1). Because the incidence of diabetes was more prevalent in female NOD mice, this study used female mice. We analyzed spontaneous diabetes incidence in mice (^+/+^, wild type; ^+/−^, heterozygous *Tyk2* KO; ^−/−^, homozygous *Tyk2* KO) in cohoused conditions after weaning to minimize the effect of different gut microbiota composition on diabetes development. As expected, the incidence of diabetes was reduced in *Tyk2^−/−^* mice compared with their *Tyk2^+/+^*and *Tyk2^+/−^* littermates (Figure 1A). Comparable diabetes development was observed between *Tyk2*^+/+^ and *Tyk2*^+/−^ mice. Although Tyk2 was reported to be associated with obesity (Derecka et al., 2012), our colony exhibited equivalent weight gain throughout the study (Figure S2). At 14 weeks of age (14w), a reduced number of islets with insulitis were observed in *Tyk2*^−/−^ mice compared with age-matched littermates, consistent with the reduction of diabetes incidence (Figures 1B and S1C). Serum anti-insulin autoantibodies (IAAs) were reduced in 14w *Tyk2*^−/−^ mice compared with age-matched *Tyk2*^+/−^ mice (Figure 1C). We also analyzed 24w *Tyk2*^−/−^ mice to assess the long-term effects of *Tyk2* deficiency. Increased numbers of inflamed islets and elevated levels of serum IAAs were observed in 24w *Tyk2*^−/−^ mice compared with 14w *Tyk2*^−/−^ mice (Figure 1B, 1C). These observations suggest that *Tyk2* deficiency does not completely prevent islet autoimmunity but reduces the progression rate of invasive insulitis leading to T1D onset.

**Figure 1.**
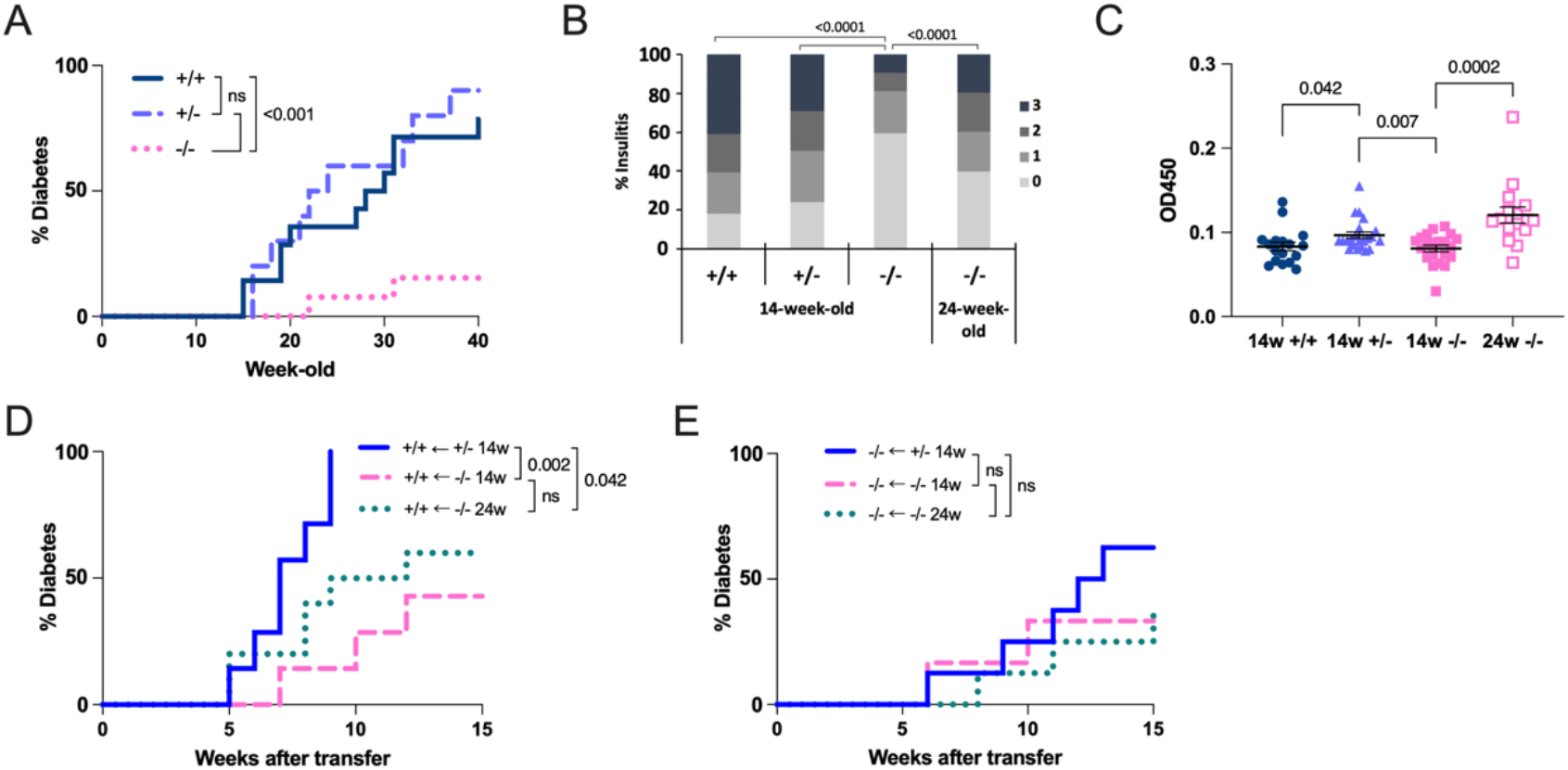
*Tyk2* deficiency suppresses autoimmune diabetes in NOD mice. (A) Incidence of spontaneous diabetes in *Tyk2*^+/+^ (n=14), *Tyk2*^+/−^ (n=10), and *Tyk2*^−/−^ NOD mice (n=13). (B) Insulitis scores of 14w (*Tyk2*^+/+^, n=8; *Tyk2*^+/−^, n=8; *Tyk2*^−/−^, n=8) and 24w *Tyk2*^−/−^ NOD mice (n=10). (C) Insulin autoantibody levels in the serum of 14w (*Tyk2*^+/+^, n=17; *Tyk2*^+/−^, n=22; *Tyk2*^−/−^, n=21) and 24w *Tyk2*^−/−^ NOD (n=16) mice. (D) Incidence of diabetes in wild type NOD.SCID mice adoptively transferred with splenocytes from 14w *Tyk2*^+/−^ NOD mice (n=7), 14w *Tyk2*^−/−^ NOD mice (n=7), or 24w *Tyk2*^−/−^ NOD mice (n=10). (E) Incidence of diabetes in *Tyk2*^−/−^ NOD.SCID mice adoptively transferred with splenocytes from 14w *Tyk2*^+/−^ NOD mice (n=8), 14w *Tyk2*^−/−^ NOD mice (n=6), or 24w *Tyk2*^−/−^ NOD mice (n=8). *P-*values were calculated using the log-rank test (A, D, E), Mann Whitney *U-*test (B), and unpaired *t-*tests (C).

### *Tyk2* expression is important for immune cells and the host environment

We next asked whether Tyk2 expression was critical for immune cells and/or the host environment (other than immune cells) using an adaptive transfer model. Because *Tyk2*^+/+^ and *Tyk2*^+/−^ mice had equivalent diabetes incidence (Figure 1A), we used splenocytes derived from *Tyk2*^+/−^ mice as a positive control. The phenotype of the transferred splenocytes, defined by CD44 and CD62L expressions (CD44^lo^CD62L^+^, naïve; CD44^hi^CD62L^-^, effector memory; CD44^hi^CD62L^+^, central memory), were comparable (Figure S1D). NOD.SCID mice, which lack T and B cells, adoptively transferred with *Tyk2*^+/−^ splenocytes developed diabetes within nine weeks after transfer, whereas mice receiving *Tyk2*^−/−^ splenocytes had reduced diabetes incidence irrespective of the donor age (Figure 1D). Next, to determine the role of *Tyk2* in the host environment, we generated *Tyk2*-deficient NOD.SCID mice and conducted the transfer experiments. We revealed that a loss of *Tyk2* in the host environment led to a reduced incidence of diabetes induced by *Tyk2*^+/−^ splenocytes compared with wild type NOD.SCID host mice (*P*=0.003) (Figure 1E). Thus, these results suggest that *Tyk2* expression in immune cells and in the host environment contributes to the development of autoimmune T1D.

### Reduced *Fas* expression and type I IFN signatures in *Tyk2* deficient β-cells

Pancreatic β-cells are a source of islet-associated autoantigens; therefore, we focused on β-cells as an important component of the host environment. We performed transcriptome analysis of β-cells purified by FluoZin-3-AM and tetramethylrhodamine ethyl ester perchlorate (TMRE) (Jayaraman, 2011)(Mine et al., 2020a) from at least six female mice to minimize the effect of individual heterogeneity on T1D development (Figure S1E). β-cells from 6w *Tyk2*^+/+^ mice and 6w *Tyk2*^−/−^ mice had similar gene expression profiles, whereas β-cells from 11w mice had different gene expression profiles from those of 6w mice (Figure 2A). This suggests that Tyk2-mediated signaling in β-cells becomes active with age. These differences in gene expression profiles in β-cells with age were confirmed by hierarchical clustering that divided the data into two clusters by age (Figures S1E). β-cells from 11w *Tyk2*^+/+^ and *Tyk2*^−/−^ mice had an enrichment in immune-related pathways compared with 6w mice (Figures 2B and S1F). Indeed, chemokines including *C-X-C motif ligand 9* (*Cxcl9*), *Cxcl10*, and *Cxcl13*, an IFN-stimulated gene (ISG) *Gbp3*, and an adhesion molecule *Vcam1*, were highly expressed in β-cells from 11w mice compared with 6w mice irrespective of the *Tyk2* genotype (Figure 2C).

**Figure 2.**
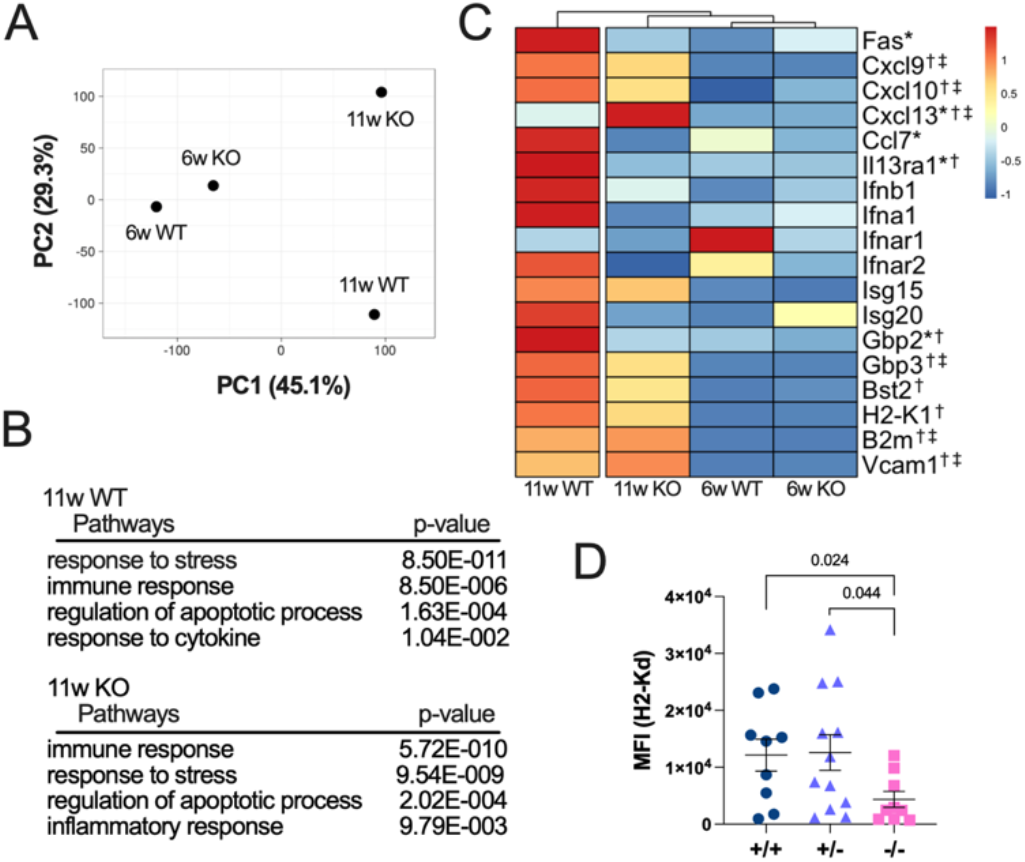
Increased inflammatory signature in β-cells with age is attenuated by *Tyk2* deficiency. (A) Principal component analysis (PCA) of transcriptome data of purified β-cells pooled from 6w *Tyk2*^+/+^ (n=6), 6w *Tyk2*^−/−^ (n=6), 11w *Tyk2*^+/+^ (n=8), and 11w *Tyk2*^−/−^ NOD mice (n=9). (B) Gene ontology (GO) biological process analysis of β-cells from 11w mice. (C) Expression heatmap of selected genes. Asterisks indicate differentially expressed genes between 11w wild type vs 11w KO mice. Daggers indicate differentially expressed genes between 6w wild type vs 11w wild type mice. Double daggers indicate differentially expressed genes between 6w KO vs 11w KO mice. (D) MHC I (H2-K^d^) mean fluorescence intensity (MFI) in β-cells from 11–12w mice (*Tyk2*^+/+^, n=9; *Tyk2*^+/−^, n=12; *Tyk2*^−/−^, n=9). *P-*values were calculated using unpaired *t-*tests.

However, *Gbp2*, an ISG, was expressed at lower levels in β-cells from 11w *Tyk2*^−/−^ mice compared with 11w *Tyk2*^+/+^ mice (Figure 2C). The ISGs including *Isg20*, *Gbp3*, and *Bst2*, and Type I IFN receptors (*Ifnar1*, *Ifnar2*) were expressed at slightly lower in β-cells from 11w *Tyk2*^−/−^ mice compared with 11w *Tyk2*^+/+^ mice. It was reported that the expressions of *Cxcl9* and *Cxcl10*, ligands for C-X-C motif chemokine receptor 3 (Cxcr3), in β-cells have a role in the chemotaxis of islet-autoreactive T cells (Uno et al., 2010). The expression levels of *Cxcl9* and *Cxcl10* in β-cells from *Tyk2*^−/−^ 11w mice were slightly reduced compared with 11w *Tyk2*^+/+^ mice. In contrast, *Cxcl13*, a ligand for Cxcr5 involved in B cell chemotaxis, was expressed at higher levels in β-cells from 11w *Tyk2*^−/−^ mice. These results suggest that type I IFN signaling in β-cells and T cell migration into inflamed islets are slightly impaired in *Tyk2*^−/−^ mice.

The increased expression of MHC I in β-cells was reported to be a feature of autoimmune T1D (Imagawa et al., 2000). *H2-K1*, an MHC I molecule, was expressed at lower levels, and *B2m*, a component of MHC I, was expressed at higher levels, in β-cells from 11w *Tyk2*^−/−^ mice compared with 11w *Tyk2*^+/+^ mice. The protein expression levels of MHC I in β-cells were decreased in 11–12w *Tyk2*^−/−^ mice compared with age-matched littermates (Figure 2D). Notably, the expression of *Fas* (*Cd95*), a cell surface death receptor involved in β-cell death (Moriwaki et al., 1999), was lower in β-cells from 11w *Tyk2*^−/−^ mice compared with 11w *Tyk2*^+/+^ mice. These results suggest that *Tyk2*^−/−^ β-cells exhibit reduced susceptibility to immune cell-mediated destruction compared with *Tyk2*^+/+^ β-cells. Together, these findings suggest that the increased inflammatory state of β-cells with age is attenuated by *Tyk2* deficiency, leading to the preservation of β-cells from immune cell attack.

### Immune cell profiles in the pancreas and pancreatic lymph node (PLN)

Tyk2 was reported to be involved in immune cell functions (Shimoda et al., 2000a)(Oyamada et al., 2009), and therefore we assessed immune cell profiles with age. T cell maturation in the thymus was comparable between *Tyk2*^+/−^ and *Tyk2*^−/−^ mice (Figure S2A). There was no difference in the frequency and number of splenic immune cells (CD4 T, CD8 T, γδ T, and B cells) with some exceptions (Figure S2B). Next, we assessed immune cells in the pancreas (Figure S2C). Although the frequency was equivalent between groups, the numbers of CD4 T, CD8 T, γδ T, and B cells in the pancreas were decreased in 11–14w *Tyk2*^−/−^ mice compared with *Tyk2*^+/−^ mice (Figures 3A, S2D, E, F). The phenotypes of CD4 T and CD8 T cells defined by CD44 and CD62L expression were also comparable between genotypes (Figure S3A). In addition, the frequency of IFN-γ or IL-17A-producing T cells, which are important effector cells in autoimmune T1D (Kuriya et al., 2013), was similar between genotypes (Figure S3B). Thus, there was reduced immune cell infiltration into the pancreas of *Tyk2*^−/−^ mice without cell-type specificity, and these T cells had no defective IFN-γ and IL-17A-producing capacity.

**Figure 3.**
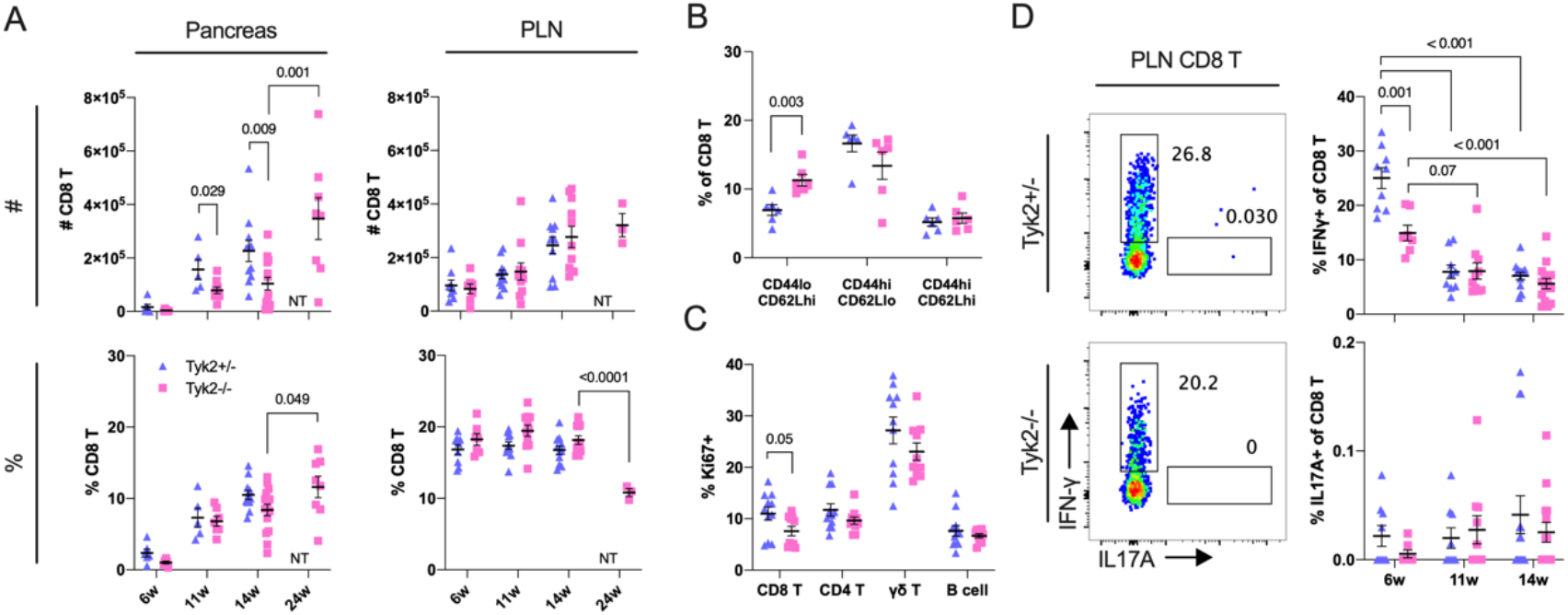
Increased naïve CD8 T cells and reduced IFN-γ-producing CD8 T cells in the PLN of *Tyk2* deficient NOD mice. (A) Cell number (#) or frequency among CD45^+^ cells (%) of CD8 T cells in the PLN and pancreas measured by flow cytometry. (B) Frequency of naïve (CD44^lo^ CD62L^+^), effector memory (CD44^hi^ CD62L^-^), and central memory (CD44^hi^ CD62L^+^) CD8 T cells in the PLN. (C) Frequency of Ki67^+^ cells among the indicated immune cells in the PLN of 6w mice. (D) Flow cytometry plots of IFN-γ and IL-17-producing CD8 T cells in the PLN of 6w mice. The graphs indicate the frequency of IFN-γ and IL-17-producing CD8 T cells among CD8 T cells in the PLN of mice at the indicated ages. *P-*values were calculated using unpaired *t-*tests.

PLN is a reservoir for islet-autoreactive T cells (Gearty et al., 2022; Gagnerault et al., 2002). Next, we analyzed the PLN and found that naïve CD8 T cells were increased in the PLN of 6w *Tyk2*^−/−^ mice compared with age-matched *Tyk2*^+/−^ mice (Figures 3B and S3C). Ki67-positive proliferating CD8 T cells were slightly decreased in the PLN of *Tyk2*^−/−^ mice compared with age-matched *Tyk2*^+/−^ mice (Figure 3C). In addition, IFN-γ-producing CD8 T cells were decreased in the PLN of 6w *Tyk2*^−/−^ mice (Figure 3D). These defects were not observed in CD8 T cells from the inguinal LN (iLN) (Figure S3D, E). There was comparable cytokine producing capacity of CD4 T cells and similar frequency of Foxp3-positive Tregs in the PLN between the genotypes (Figure S3F, G). Although these observations in CD8 T cells were restricted to the initiation phase of autoimmune T1D when the PLN was crucial for the development of diabetes (∼10w) (Gagnerault et al., 2002), these results suggest that the defective proliferation and/or reduced number of CD8^+^ CTLs, which are the primary effector immune cells of β-cell destruction (Gearty et al., 2022), in the PLN are responsible for the inhibition of autoimmune T1D development in *Tyk2*^−/−^ NOD mice.

### Reduced proliferation of CD8 T cells in the PLN of *Tyk2* deficient mice

Antigen encountered naïve T cells become activated, proliferate, and develop into effector and memory T cells (Kurts et al., 2010). To investigate the role of Tyk2 in the priming of naïve CD8 T cells, we used islet-specific glucose-6-phosphatase catalytic subunit-related protein (IGRP)-specific CD8 T cells (8.3 CD8 T cells) from NY8.3-NOD transgenic mice, which express a T cell receptor (TCR) specific for the islet autoantigen IGRP_206-214_ peptide (Verdaguer et al., 1997). The transfer of *Tyk2*^+/−^ or *Tyk2*^−/−^ naïve 8.3 CD8 T cells into 6–8w *Tyk2*^+/−^ recipient mice resulted in a similar upregulation of CD44, a marker of an antigen experienced phenotype, in the transferred cells, and proliferation in the PLN 5d post transfer, whereas their proliferation was suppressed in the PLN of age-matched *Tyk2*^−/−^ recipient mice (Figures 4A and S4A). Ten days after the transfer, *Tyk2*^+/−^ naïve 8.3 CD8 T cells showed greater proliferation in the PLN of *Tyk2*^−/−^ recipient mice compared with 5 days post transfer (Figure 4A). The limited proliferation of 8.3 CD8 T cells was observed in the iLN (Figure S4B), indicating the importance of the PLN for the proliferation of islet-autoreactive CTLs. Together, these observations suggest that CD8 T cell-extrinsic mechanisms have a role in the proliferation of islet-autoreactive CTLs in the PLN.

**Figure 4.**
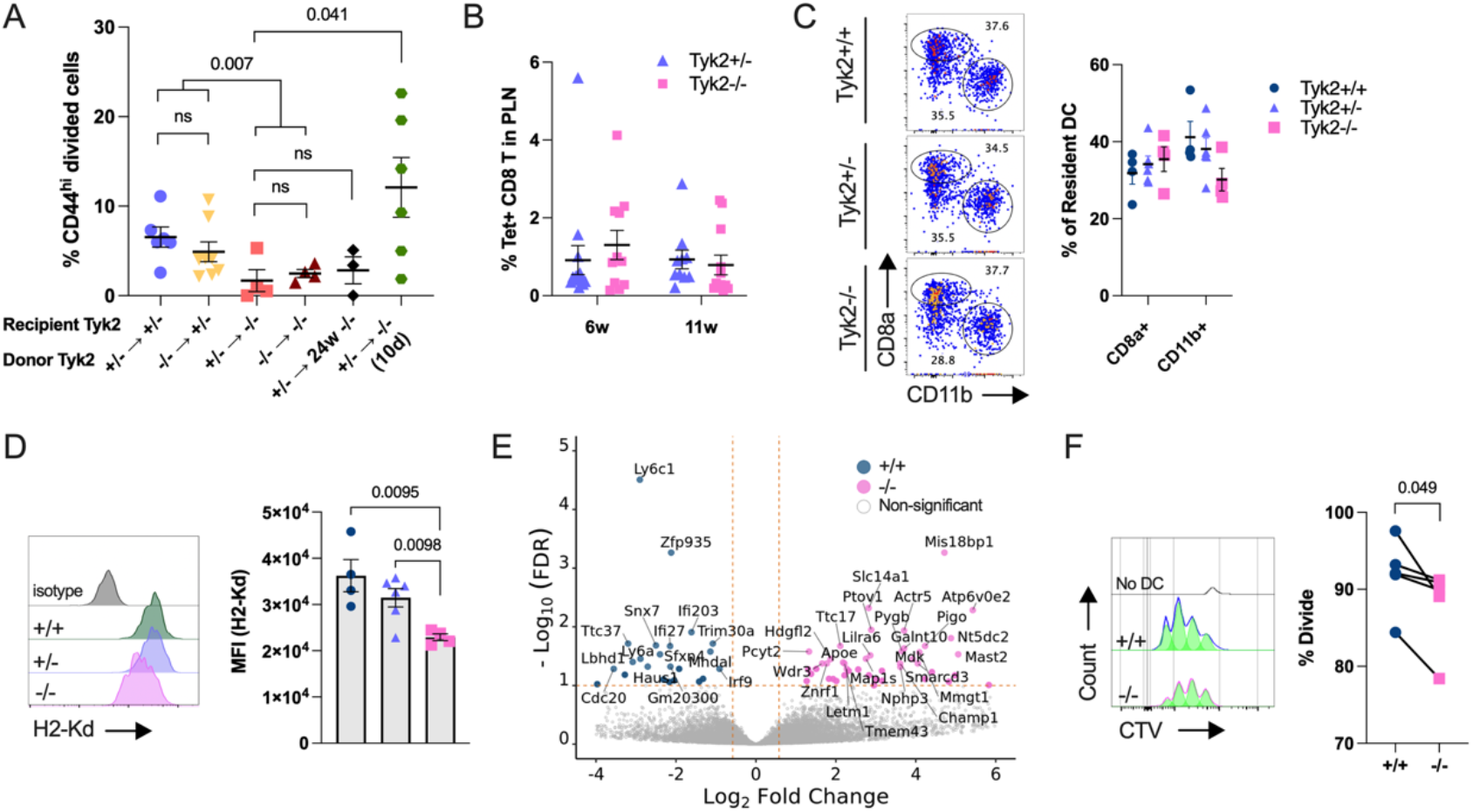
Defective APC function of CD8^+^ rDC in *Tyk2* deficient NOD mice. (A) Frequency of proliferating *Tyk2*^+/−^ or *Tyk2*^−/−^ CTV-labeled 8.3 CD8 T cells in the PLN of the indicated recipient mice. (B) Frequency of IGRP peptide-specific CD8 T cells in the PLN. (C) Flow cytometry plots of resident DC (rDC) in the PLN of 6w mice. The frequencies of CD8^+^ rDC and CD11b^+^ rDC among rDC are shown (*Tyk2*^+/+^, n=4; *Tyk2*^+/−^, n=6; *Tyk2*^−/−^, n=4). (D) Expression of H2-K^d^ in CD8^+^ rDC in the PLN of 6w mice (*Tyk2*^+/+^, n=4; *Tyk2*^+/−^, n=6; *Tyk2*^−/−^, n=4). (E) Volcano plot of CD8^+^ rDC in the PLN of 6w mice (*Tyk2*^+/+^, n=4; *Tyk2*^−/−^, n=4). (F) Proliferation of 8.3 CD8 T cells labeled with CTV and co-cultured with CD8^+^ rDC for 3d.

Because IGRP epitopes were reported to appear and spread in the middle-to-late stage of T1D (Luo et al., 2010), defects in IGRP epitope spreading in *Tyk2*^−/−^ mice might correlate with the reduced proliferation of CD8 T cells in the PLN of *Tyk2*^−/−^ mice. To exclude this possibility, we evaluated IGRP-specific CD8 T cells (IGRP CD8 T) using tetramers containing a mimotope of IGRP (NRP-V7). A comparable frequency of IGRP CD8 T cells was observed in the PLN of both *Tyk2* genotypes (Figure 4B). Furthermore, the proliferation of 8.3 CD8 T cells was also suppressed in 24w *Tyk2*^−/−^ recipient mice that had advanced insulitis compared with 14w *Tyk2*^−/−^ mice (Figures 1B and 4A). These data indicate that the defective proliferation of CD8 T cells in the PLN of *Tyk2*^−/−^ mice is not related to reduced IGRP epitope spreading in *Tyk2*^−/−^ mice.

### MHC I expression is reduced in *Tyk2* deficient CD8^+^ resident DCs

The results of the adaptive transfer experiments (Figure 4A) suggest that *Tyk2* expression in antigen presenting cells (APCs) was associated with the efficient proliferation of islet-autoreactive CTLs in the PLN. A previous study reported the importance of Tyk2 in antigen presentation by dendritic cells (DCs) for the expansion of IFN-γ^+^ CD8 T cells following *Listeria monocytogenes* infection (Aizu et al., 2006). Next, we analyzed DCs, which have a central role in the adaptive immune system via antigen presentation to T cells (Figure S4C) (Kurts et al., 2010). In the PLN, the frequency of CD11c^+^ MHC II^mid^ resident DCs (rDCs), including CD8^+^ and CD11b^+^ subsets, and CD11c^+^ MHC II^hi^ migratory DCs (mDCs), including the CD103^+^ CD11b^-^, CD103^+^ CD11b^+^, and CD103^-^ CD11b^+^ subsets, were comparable irrespective of *Tyk2* genotype (Figures 4C and Figure S4D). Comparable expression levels of MHC II (I-A^g7^) were observed in these DC subsets from all genotypes (Figure S4E). However, we found that *Tyk2* deficiency reduced the expression of MHC I (H2-K^d^), a crucial molecule for presenting antigens to CD8 T cells (Kurts et al., 2010), in CD8^+^ rDC but not other subsets of DCs, except for CD103^-^ CD11b^+^ mDC (Figures 4D and S4F). This MHC I reduction in CD8^+^ rDC was not specific for the PLN as comparable results were observed in the iLN and spleen (Figure S4G). The expression levels of co-stimulatory molecules including CD40 and CD86 in CD8^+^ rDC were comparable between genotypes (Figure S4H). Previous studies reported that CD8^+^ rDC have a role in the expansion of CTLs by cross-priming (Kurts et al., 2010; Hildner et al., 2008) and therefore we focused on CD8^+^ rDC. We analyzed the gene expression profiles of CD8^+^ rDC in the PLN and found that *Irf9* and ISGs including *Ifi27* and *Ifi203* were highly expressed in *Tyk2*^+/+^ CD8^+^ rDC compared with *Tyk2*^−/−^ cells (Figure 4E), suggesting that type I IFN signaling is upregulated in *Tyk2*^+/+^ CD8^+^ rDC (Figure S4I). The expression of IL-12 and IL-23, which are involved in T cell differentiation via Tyk2, were comparable between genotypes. To assess the APC function of CD8^+^ rDC, we performed an *ex vivo* co-culture experiment. This revealed that *Tyk2*^−/−^ CD8^+^ rDC induced lower levels of 8.3 CD8 T cells expansion compared with *Tyk2*^+/−^ CD8^+^ rDC (Figure 4F). These results suggest defective Tyk2-mediated type I IFN signaling in CD8^+^ rDC results in decreased MHC I expression, leading to the impaired cross-priming of islet-autoreactive CTLs.

### Reduced T-bet^+^ CD8 T cells in the PLN of *Tyk2* deficient mice

Although we found *Tyk2*^−/−^ CD8^+^ rDC had defective APC functions, the number of CD8 T cells and IGRP CD8 T cells in the PLN were comparable between the *Tyk2* genotypes (Figures 3A, 4B). Therefore, we next characterized antigen experienced CD8 T cells in the PLN. Because transcription factors (TFs) are key regulators of T cell functions, we examined TFs in CD8 T cells. The frequency of Eomes, Blimp-1, and Tcf1-positive CD44^hi^ polyclonal or IGRP CD8 T cells in the PLN were comparable between genotypes (Figure S4J, K). We found that polyclonal and IGRP-specific CD44^hi^ CD8 T cells positive for T-bet, an important TF for the production of IFN-γ, were decreased in the PLN of *Tyk2*^−/−^ mice compared with *Tyk2* sufficient mice (Figure 5A, B), supporting the results of *ex vivo* cytokine-producing experiments (Figure 3D). The expression levels of T-bet were also reduced in CD44^hi^ T-bet^+^ IGRP CD8 T cells in the PLN of Tyk2^−/−^ mice compared with *Tyk2*^+/−^ mice (Figure 5C). Thus, these data suggest that the development of T-bet^+^ CTLs is impaired in the PLN of *Tyk2*^−/−^ NOD mice. IL-12 is known to drive T-bet expression in T cells during priming (Starbeck-Miller et al., 2014). Next, to address the role of *Tyk2* in CD8 T cells during priming, purified CD8 T cells were stimulated *ex vivo* with anti-CD3/CD28 beads in the presence of IL-2 and IL-12. We found that the loss of *Tyk2* in CD8 T cells was not associated with the expansion of T-bet^+^ CD8 T cells (Figure 5D). However, the IL-12-induced upregulation of T-bet expression levels was impaired in *Tyk2*^−/−^ CD8 T cells compared with *Tyk2*^+/−^ cells (Figure 5E). These data suggest that *Tyk2*^−/−^ CD8 T cells develop CTLs that are expressed lower levels of T-bet than those of *Tyk2* sufficient CD8 T cells in the PLN due to defective IL-12 signaling (Figure 5C).

**Figure 5.**
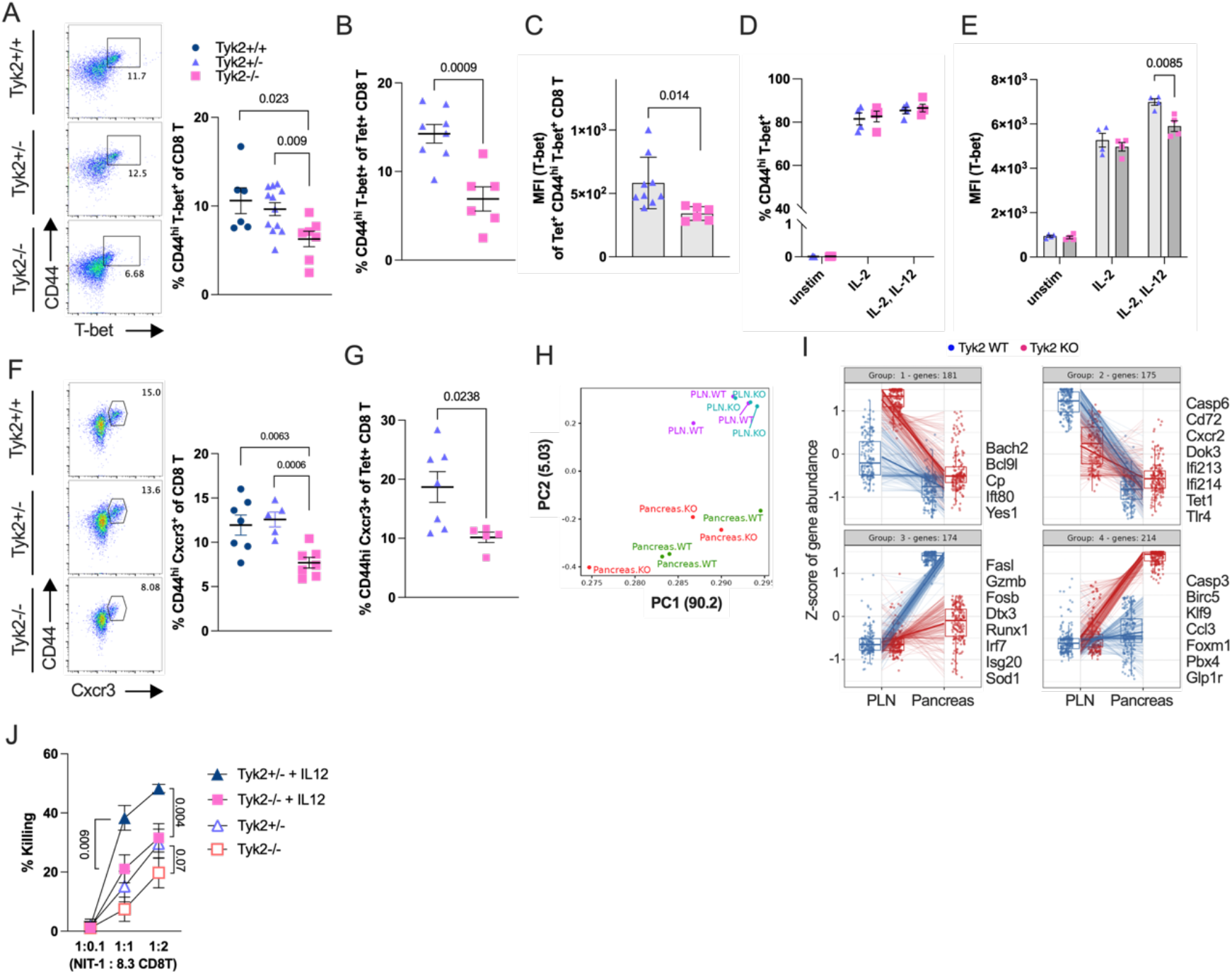
Defective development of T-bet^+^ CTLs in the PLN and effector function of CTLs in the pancreas of *Tyk2* deficient NOD mice. (A) Flow cytometry plots and graph indicate the frequency of CD44^hi^ T-bet^+^ polyclonal cells among CD8 T cells in the PLN of 6w mice. (B) Frequency of CD44^hi^ T-bet^+^ cells among IGRP-specific CD8 T cells in the PLN of 6w mice. (C) Expression of T-bet in CD44^hi^ T-bet^+^ IGRP-specific CD8 T cells in the PLN of 6w mice. (D) Frequency of CD44^hi^ T-bet^+^ cells in *ex vivo*-activated *Tyk2*^+/−^ or *Tyk2*^−/−^ CD8 T cells with or without IL-12 for 3d. (E) Expression of T-bet in *ex vivo*-activated *Tyk2*^+/−^ or *Tyk2*^−/−^ CD8 T cells with or without IL-12 for 3d. (F) Flow cytometry plots and graph indicate the frequency of CD44^hi^ Cxcr3^+^ cells among polyclonal CD8 T cells in the PLN of 6w mice. (G) Frequency of CD44^hi^ Cxcr3^+^ cells among IGRP-specific CD8 T cells in the PLN of 6w mice. (H) PCA of transcriptome data of purified CD44^hi^ IGRP-specific CD8 T cells in the PLN (*Tyk2*^+/+^, n=3, *Tyk2*^−/−^, n=3) and IGRP-specific CD8 T cells in the pancreas (*Tyk2*^+/+^, n=3, *Tyk2*^−/−^, n=3). (I) Box plots show the pattern of differentially expressed genes. Selected gene names were highlighted. (J) The killing capacity of *ex vivo*-activated 8.3 CD8 T cells with or without IL-12 against NIT-1 cells at the indicated ratios (NIT-1:8.3 CD8 T). The data represent two independent experiments with similar results.

Chemotaxis is an important mechanism in the induction of inflammation. It was reported that a target gene of T-bet is *Cxcr3* (Lazarevic et al., 2013), a receptor for Cxcl9 and Cxcl10, which were increased in β-cells with age (Figure 2C). Next, we examined the expression of Cxcr3 and found that polyclonal and IGRP-specific CD44^hi^ Cxcr3^+^ CD8 T cells were decreased in the PLN of *Tyk2*^−/−^ mice compared with age-matched littermates (Figures 5F, 5G and S4L). The decreased number of CD44^hi^ Cxcr3^+^ CD8 T cells was specific for the PLN of *Tyk2*^−/−^ mice because there were no differences in the iLN and spleen (Figure S4M). In contrast, Cxcr3 expression in CD4 T cells in the PLN was comparable between genotypes (Figure S4L). Together, these observations suggest that the comparable number of CD8 T cells in the PLN between *Tyk2* genotypes (Figure 3A) might be related to the developed CTLs expressing Cxcr3, exiting the PLN, and migrating to the islets, resulting in similar numbers of CD8 T cells in PLN but increased numbers of CD8 T cells in the pancreas of *Tyk2* sufficient mice compared with *Tyk2*^−/−^ mice (Figure 3A).

### Cytotoxic functions of *Tyk2* deficient CTLs are impaired

Our data suggest that the CD8^+^ rDC-mediated cross-priming of CD8 T cells and the development of islet-autoreactive T-bet^+^ CTLs were impaired but not abolished in the PLN of *Tyk2*^−/−^ mice. It was demonstrated that stem-like CD8 T cells in the PLN give rise to pathogenic CD8 T cells in the pancreas with upregulating effector functions (Gearty et al., 2022). Next, to test whether the transition of gene expression profiles in autoreactive CTLs depends on *Tyk2* genotype, we performed the transcriptome analysis of purified antigen-experienced CD44^hi^ IGRP CD8 T cells in the PLN and IGRP CD8 T cells in the pancreas (Figure 5H). The gene expression profiles of IGRP-specific CD8 T cells in the pancreas differed from those in the PLN (Gearty et al., 2022) irrespective of the *Tyk2* genotype (Figure 5H). We found that 744 genes were differentially expressed and that *Fasl*, *Gzmb*, and ISGs including *Irf7* and *Isg20* were upregulated in *Tyk2*^+/+^ IGRP CD8 T cells in the pancreas compared with *Tyk2*^−/−^ cells (Group 3; Figure 5I). In contrast, *Tyk2*^−/−^ IGRP CD8 T cells purified from the pancreas upregulated apoptosis-related genes including *Casp3*and *Birc5* (Group 4; Figure 5I). These results suggest that the loss of *Tyk2* partially impairs the effector functions of islet-autoreactive CTLs.

To assess the cytotoxic activity of CTLs, *ex vivo*-activated 8.3 CD8 T cells (8.3 CTLs) were cultured with NIT-1 cells, a pancreatic β-cell line derived from NOD mice (Hamaguchi et al., 1991), and the survival of NIT-1 cells was analyzed (Zhang et al., 2014). The cytotoxic activity of *Tyk2*^−/−^ and *Tyk2*^+/−^ 8.3 CTLs was comparable when 8.3 CD8 T cells were activated without IL-12; however, *Tyk2*^−/−^ 8.3 CTLs stimulated with IL-12 had a reduced killing capacity compared with those of *Tyk2*^+/−^ 8.3 CTLs (Figure 5J). These results suggest that defects in endogenous IL-12 signaling in CTLs associated with *Tyk2* deficiency reduces the cytotoxic activity of CTLs against β-cells.

### Tyk2 inhibitor has the potential to prevent the development of autoimmune T1D

Recently, a selective Tyk2 inhibitor, BMS-986165 (Wrobleski et al., 2019) was reported to inhibit autoimmune diseases including psoriasis in mice and humans (Burke et al., 2019; Papp et al., 2018), although its effects on autoimmune T1D are unknown. Our results suggest that Tyk2 is a potential target for preventing the development of autoimmune T1D. To test whether BMS-986165 inhibited the development of CTLs, purified CD8 T cells were stimulated *ex vivo* with anti-CD3/CD28 beads, IL-2, and IL-12, in the presence or absence of BMS-986165. The expression levels of T-bet in stimulated CD8 T cells were reduced dose-dependently (Figure 6A). Consistent with the T-bet expression levels, the β-cell killing capacity of stimulated 8.3 CD8 T cells was impaired dose-dependently (Figure 6B), supporting the results with Tyk2^−/−^ 8.3 CD8 T cells (Figure 5J). Next, we assessed Cxcr3 expression levels in stimulated CD8 T cells. Re-culture of stimulated CD8 T cells for 2 days induced Cxcr3 expression in the cells (Figure 6C) (Collins et al., 2002). However, re-cultured CD8 T cells primed with BMS-986165 had reduced Cxcr3 expression levels dose-dependently (Figure 6C), indicating that the reduction of Cxcr3 in CTLs was due to a CTL intrinsic mechanism. Next, to test the effect of BMS-986165 on developed CTLs, we treated stimulated CD8 T cells with BMS-986165 for 2 days and found that the expression levels of T-bet and Cxcr3 were unchanged (Figure S5A, B). However, BMS-986165 inhibited IL-12 and IL-18-induced bystander IFN-γ production in CD44^hi^ CD8 T cells dose-dependently (Figure S5B). These results suggest that Tyk2 has a role in the development of T-bet^+^ Cxcr3^+^ CTLs and bystander activation, and is dispensable for the maintenance of T-bet and Cxcr3 expression in CTLs.

**Figure 6.**
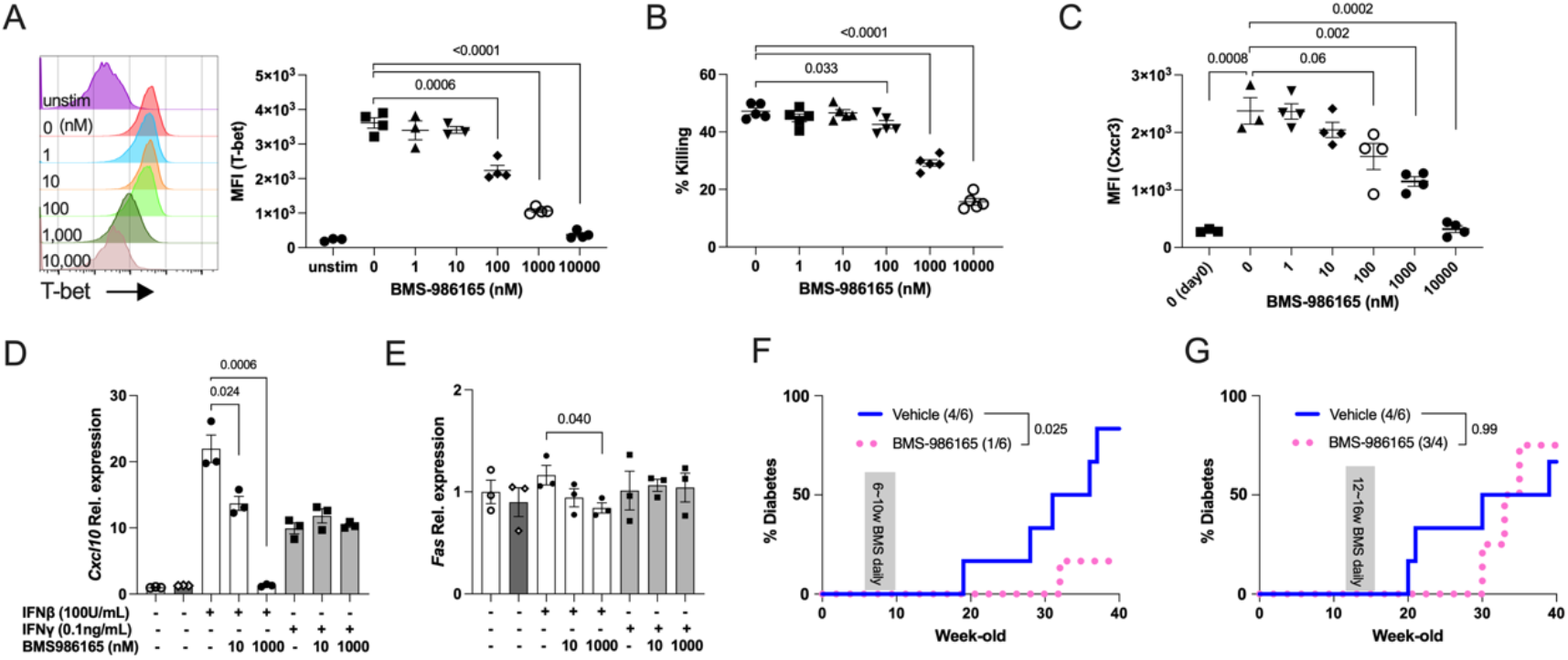
BMS-986165 prevents the development of autoimmune T1D. (A) Expression of T-bet in stimulated CD8 T cells in the presence of BMS-986165. (B) The β-cell killing capacity of *ex vivo*-activated 8.3 CD8 T cells in the presence of BMS-986165. (C) Expression of Cxcr3 in stimulated CD8 T cells with BMS-986165. *Ex vivo* activated CD8 T cells were re-cultured without BMS-986165 for 48 hours and the induction of Cxcr3 expression was analyzed. Day 0 data indicate the Cxcr3 expression levels just after stimulation. (D) Expression of *Cxcl10* in NIT-1 cells stimulated with IFN-β or IFN-γ in the presence or absence of BMS-986165 for 3 hours. (E) Expression levels of *Fas* in NIT-1 cells stimulated with IFN-β or INF-γ in the presence or absence of BMS-986165 for 24 hours. The data represent two independent experiments with similar results. (F) Incidence of diabetes in vehicle (n=6) or BMS-986165-treated (n=6) wild type NOD mice for 4 weeks from 6w of age. (G) Incidence of diabetes in vehicle (n=6) or BMS-986165-treated (n=4) wild type NOD mice for 4 weeks from 12w of age. *P-*values were calculated using unpaired *t-*tests (A-E) and log-rank test (F, G).

Next, to test whether BMS-986165 inhibited inflammatory responses in β-cells, NIT-1 cells with or without BMS-986165 were stimulated with type I or type II IFNs. BMS-986165 inhibited IFN-β-induced *Cxcl9*, *Cxcl10*, and *Isg15* expressions dose-dependently (Figures 6D and S5D, E). BMS-986165 had no effect on IFN-γ-induced *Cxcl9, Cxcl10*, and *Isg15* expressions in NIT-1 cells, because Tyk2 is not associated with type II IFN signaling mediated by other Jaks, confirming the high selectively of the Tyk2 inhibitor. We found that IFN-β-induced *Fas* expression was reduced by BMS-986165 dose-dependently. Together, these results demonstrate that BMS-986165 inhibits the development and bystander activation of CTLs, and type I IFN-induced gene expressions in β-cells.

To demonstrate the effects of BMS-986165 on the development of autoimmune T1D *in vivo*, we administrated the inhibitor to NOD mice by oral gavage once-daily at 30 mg/kg for 4 weeks from 6w of age. NOD mice treated with BMS-986165 had reduced diabetes incidence (1/6) compared with vehicle control mice (5/6) (*p*=0.025, Figure 6F). In contrast, treatment with BMS-986165 at the same dose from 12w of age resulted in comparable diabetes incidence (3/4) with vehicle control mice (4/6) (*p*=0.99, Figure 6G). These results suggest that treatment with a Tyk2 inhibitor started at an early stage of T1D may be a potential prevention strategy to prevent the development of autoimmune T1D.

## DISCUSSION

Here, we demonstrate that the blockade of Tyk2-mediated signaling in autoimmune T1D model NOD mice impaired the development and effector function of CTLs, efficient cross-priming of CTLs by CD8^+^ rDC, and expression of inflammatory molecules in β-cells. In the context of autoimmune disease, the role of Tyk2 in CD4 T cells, but not CD8 T cells, has been well described (Oyamada et al., 2009; Muromoto et al., 2021). Our study reveals that IL-12 signaling via Tyk2 upregulated T-bet expression in CTLs during their priming, leading to the upregulation of Cxcr3 and induction of effector functions against β-cells. In addition, the efficient cross-priming of CTLs by CD8^+^ rDC may be associated with MHC I expression levels in CD8^+^ rDC, probably induced by type I IFN signaling via Tyk2. These data provide evidence that Tyk2-mediated signaling has a critical role in the generation of autoreactive CD8^+^ CTLs. However, increased naïve CD8 T cells and decreased IFN-γ-producing CD8 T cells in the PLN of *Tyk2*^−/−^ mice were limited in the initiation phase of T1D (Figure 1B, D) when the PLN is dispensable for T1D development (Gagnerault et al., 2002). This is likely because loss or inhibition of Tyk2 did not completely inhibit CTL priming (Figures 4A, 5, 6B), the number of CTLs in the PLN became comparable to that in the PLN of *Tyk2*-sufficient mice in accordance with aging. Although the development of CTLs in the PLN of *Tyk2*^−/−^ mice was not abolished, the reduced killing capacity of *Tyk2*^−/−^ CTLs against β-cells probably contributes to the suppression of the development of autoimmune T1D. Consequently, our study reveals that Tyk2 may be a good therapeutic target for not only CD4 T cell-related but also CD8 T cell-related autoimmune diseases.

Consistent with an earlier study (Ishizaki et al., 2011), a similar frequency of Treg was observed regardless of the *Tyk2* genotype (Figure S3G). The non-involvement of Tyk2 in Treg differentiation and maintenance (Ishizaki et al., 2011) may have led to the efficient suppression of CTLs in *Tyk2*^−/−^ mice. Although the differentiation of Th1 and Th17 cells in the PLN was comparable between *Tyk2* genotypes, bystander IFN-γ production by CD44^hi^ CD4 T cells was impaired by Tyk2 inhibition (Figure S5C), suggesting the pathogenic role of islet-autoreactive CD4 T cells may be also defective in *Tyk2*^−/−^ mice.

Our data suggest that Tyk2-mediated signaling in β-cells becomes active with age. Because the expression levels of chemokines, ISGs, and MHC I molecules were lower in the β-cells from 6w mice compared with those from 11w mice (Figure 2C, D), the increased inflammatory signature of β-cells with age may be caused by infiltrating CTLs or the local islet environment induced by CTLs. We found that *Cxcl9* and *Cxcl10* expressions in β-cells were upregulated with age, which were slightly attenuated by *Tyk2* deficiency, at least in part, due to impaired type I IFN signaling in β-cells (Figure 6D and S5D). In addition to the low *Cxcl9* and *Cxcl10* expressions, reduced MHC I and *Fas* expressions in the β-cells of *Tyk2*^−/−^ mice may contribute to the preservation of β-cells from CTL attack. We analyzed purified β-cells using the TMRE dye (Jayaraman, 2011), a marker of mitochondrial membrane potential, which is rapidly decreased in apoptotic cells (Kuznetsov et al., 2011). Although the apoptosis-related pathway was enriched in β-cells from 11w mice (Figure 2B), our transcriptome datasets of β-cells may have excluded some β-cells that had started to undergo cell death by apoptosis. Given that apoptotic cells release chemotactic factors such as Ccl2 and IL-8 (Cullen et al., 2013), β-cells undergoing apoptosis may have a role in the development of insulitis via mechanisms different from live β-cells that express *Cxcl9* and *Cxcl10*.

Together, these findings suggested that the mechanisms described above act in concert to inhibit the development of autoimmune T1D in *Tyk2*^−/−^ NOD mice. In support of this, a recent study reported that the loss-of-function variant rs34536443 in *TYK2* was associated with a lower risk of autoimmune disease including T1D (Yuan et al., 2023). Thus, Tyk2 is involved in the pathogenesis of autoimmune T1D in mice and humans.

Treatment with the Tyk2 inhibitor BMS-986165 impaired the development of T-bet^+^ Cxcr3^+^ CTLs, bystander activation of CTLs, and *Fas,* chemokine, and ISG expressions in β-cells dose-dependently. A reduced T1D progression rate was evident when the treatment was started at 6w but not at 12w of age, suggesting that the initiation or early phase of T1D is the therapeutic window for the Tyk2 inhibitor. We found that IFN-γ-producing CD8 T cells in the PLN were more abundant in the initiation phase of T1D than in the late phase (Figure 3D), indicating the initiation phase of T1D is optimal for the efficient suppression of CTL development in the PLN. Because CD8 T cells are the main effector immune cells in the development of T1D (Gearty et al., 2022), early intervention and longitudinal treatment with a Tyk2 inhibitor may be a potent immunotherapeutic strategy for autoimmune T1D.

Tyk2 is involved in anti-microorganism responses via cytokine signaling (Izumi et al., 2015; Kreins et al., 2015; Aizu et al., 2006; Shimoda et al., 2000b). Clinical studies reported potential adverse effects of BMS-986165, including nasopharyngitis, diarrhea, nausea, and upper respiratory tract infection (Strober et al., 2023; Yuan et al., 2023). In a clinical trial, BMS-986165 was considered safe on the basis of observations of a slight increase but low rate of serious viral infection, probably due to the high selectivity of the Tyk2 inhibitor over other Jaks, resulting in reduced but not abolished antiviral responses in the patients (Strober et al., 2023). However, it was reported that some patients with T1D, especially with acute onset and fulminant T1D, exhibited flu-like syndromes at disease onset or elevated antibody levels against viruses in the blood (Imagawa et al., 2003; Nagafuchi et al., 2015; Hosokawa et al., 2019). Some presenting symptoms were non-serious such as fever, cough, and sore throat (Nagafuchi et al., 2015; Imagawa et al., 2003) although some severe cases have been reported (Vreugdenhil et al., 2000; Müller et al., 2021)(Jenson et al., 1980). In contrast to the role of *Tyk2* in autoimmune T1D described in this paper, we reported that reduced *Tyk2* expression was a risk for virus-induced diabetes and impaired insulin secretion (Izumi et al., 2015; Nagafuchi et al., 2015; Mine et al., 2017; Mori et al., 2021). The importance of anti-viral defense in β-cells to prevent β-cell destruction caused by viruses was reported based on *in vivo* findings (Flodström et al., 2002; Izumi et al., 2015; Mine et al., 2020a; Lalwani et al., 2019). Although the role of viruses in T1D is controversial, the advantages and risks of immunomodulation (Gorman et al., 2017) should be considered when developing treatments for T1D. An anti-diabetogenic virus vaccine, which is currently under development (Stone et al., 2020), would be a potent option to eliminate the potential viral contribution to β-cell destruction in patients taking immunosuppressant drugs such as Tyk2 inhibitors and anti-CD3 antibodies (Herold et al., 2019).

In conclusion, our study demonstrated the role of *Tyk2* in the development of autoreactive CTLs and pathogenesis of autoimmune T1D, indicating TYK2 is a potential therapeutic target for autoimmune T1D. The evaluation of candidate genes by multifaceted approaches will aid our understanding of the complex role of genes involved in multifactorial diseases and help the development of the safe and effective preventive/therapeutic strategies.

## MATERIALS AND METHODS

### Animals

Wild type NOD/shiJcl mice were obtained from Clea Japan, Inc. (Tokyo, Japan) and bred in our facility. *Tyk2* knockout mice on a C57BL/6J genetic background were generated as previously described (Shimoda et al., 2000a)(Izumi et al., 2015). *Tyk2*KO mice on a NOD/shiJcl genetic background (*Tyk2*KO.NOD) were generated by backcrossing with NOD/shiJcl mice more than ten times. All microsatellite repeat polymorphisms at the Idd loci were tested by PCR as previously described (Kim et al., 2021) (Figure S1A). *Tyk2* heterozygous and homozygote mice were healthy and fertile. NOD.SCID mice were obtained from the Jackson Laboratory Japan, Inc. (Yokohama, Japan) and bred in our facility. *Tyk2*KO mice on a NOD/SCID genetic background (*Tyk2*KO.NOD.SCID) were generated by crossing with *Tyk2*KO.NOD mice. The *Prkdcscid* gene mutation was determined by PCR as previously described (Maruyama et al., 2002). NY8.3 NOD mice were provided by S. Akazawa and N. Abiru (Nagasaki University, Nagasaki, Japan) (RRID:IMSR_JAX:005868). *Tyk2*KO mice on a NY8.3 NOD genetic background (*Tyk2*KO 8.3 NOD) were generated by crossing with *Tyk2*KO.NOD mice. Blood glucose levels were measured in tail vein blood using Glutest Ai (Sanwa, Nagoya, Japan). Diabetes was defined by a non-fasting blood glucose level exceeding 250 mg/dL. All mice were maintained on a 12-hour light/dark cycle in a temperature-controlled facility under specific pathogen-free conditions at 23±2℃ with 50±10% humidity, and provided with sterile food and water *ad libitum*. SCID mice were fed with sterile water. This study was approved by the Saga University Animal Care and Use Committee and conducted in accordance with the regulations on animal experimentation at Saga University.

### Histology

Pancreas specimens were immersed in 10% (w/v) formaldehyde overnight at 4℃ and then embedded in paraffin. Cut sections (3 µm) were subjected to hematoxylin and eosin (H&E) staining. Deparaffinization of paraffin-embedded sections was performed as follows: incubation in xylene for 5 min at room temperature (RT) ×3, 100% ethanol for 5 min at RT ×2, 90% ethanol for 5 min at RT ×1, 80% ethanol for 5 min at RT ×1, 80% ethanol for 5 min at RT ×1, and ultrapure water for 5 min at RT ×1.

### ELISA

IAA levels in serum were measured using an ELISA kit (Cusabio, Huston, TX, USA) according to the manufacturer’s instructions. The optical density was determined at 450 nm.

### Adoptive transfer experiments

Donor splenocytes were purified from the spleens of non-diabetic 14w or 24w female mice. Splenocytes (1×10^7^) were transferred into 6w to 8w wild type or *Tyk2*KO. NOD.SCID recipient mice. Blood glucose levels were measured in tail vein blood once weekly using Glutest Ai (Sanwa). To analyze the proliferation of CD8 T cells in LNs, naïve 8.3 CD8 T cells from the LNs of non-diabetic NY8.3 NOD mice were purified using two rounds of the naïve CD8 T cell isolation kit (Miltenyi Biotec, Bergisch Gladbach, Germany) with LS columns (Miltenyi Biotec). The average purity of the sorted cells was 96%. Purified naïve 8.3 CD8 T cells were labeled using a CellTrace violet cell proliferation kit (Invitrogen, Waltham, MA, USA) according to the manufacturer’s protocol. Then, 4×10^6^ labeled naïve 8.3 CD8 T cells were transferred into 6w to 9w recipient mice. Next, 5 or 10d after the transfer, the PLN and iLN were harvested and the proliferation of CTV labeled cells was analyzed using a FACSVerse flow cytometer (BD Biosciences).

### Islets isolation and β-cell sorting

Islets were isolated using 1 mg/ml collagenase-P (Roche) and Histopaque-1077 (Sigma-Aldrich, Tokyo, Japan) as previously described (Zmuda et al., 2011) with minor modifications. Purification of β-cells was performed as previously described (Jayaraman, 2011)(Mine et al., 2020a). Briefly, isolated islets were dissociated using TrypLE express (Gibco BRL, Grand Island, NY, USA) for 9 min at 37 ℃ with vortexing every 3 minutes. Dissociated islet cells were washed with RPMI-1640 containing 10% FBS and 1% PcSM, and resuspended in the medium. Dissociated islet cells were stained with FluoZin-3-AM (Invitrogen), TMRE (Life Technologies, Carlsbad, CA, USA), anti-Cd45 antibody (BioLegend, San Diego, CA, USA), and anti-H2-Kd (MHC class I) antibody (BioLegend) for 30 min at 37°C. Then, 1 mg/mL propidium iodide (Sigma-Aldrich) was added to the cell suspension just before sorting to exclude dead cells. FluoZin-3-AM-positive, TMRE-positive, and CD45-negative cells were sorted using MA900 (Sony, Tokyo, Japan), and data were analyzed using FlowJo software (Tree Star, Ashland, OR, USA).

### Microarray

We used an Affymetrix Clariom D Assay Mouse Array for the β-cell transcriptome analysis. Microarray analysis was provided by Cell Innovator (Fukuoka, Japan). To identify differentially-expressed genes, we calculated the Z scores and ratios (non-log-scale fold-change) from the normalized signal intensities for each probe to compare control and experimental samples. Criteria used for identifying up- and downregulated genes were as follows: upregulated genes, Z score > 2.0 and ratio < 1.5; downregulated genes, Z score < −2.0 and ratio > 0.66. The accession number for the data is GEO: GSE235670. Microarray analysis was supported by Cell Innovator (Fukuoka, Japan).

### Flow Cytometry Analysis

To isolate immune cells from the LNs, thymus, or spleen, tissues were minced and enzymatically dissociated with 1 mg/mL collagenase D (Roche Applied Science) and 25 U/mL DNase I (Fujifilm Wako, Osaka, Japan) in DMEM (Fujifilm Wako) for 30 min at 37°C. The digested tissues were filtered through a 70-µm cell strainer with a syringe plunger. Red blood cell lysis was performed by incubation with ammonium-chloride-potassium lysis buffer for 4 min on ice. Isolated cells were preincubated with CD16/32 blocker (clone 93; BioLegend) for 5 minutes at 4°C to prevent nonspecific staining, and stained for 20 min at 4°C with antibodies. We added 1 mg/mL propidium iodide to the cell suspension to exclude dead cells.

To isolate immune cells from the pancreas, pancreases were perfused with 1 mg/mL collagenase D and 0.5 mg/mL Trypsin inhibitor (Fujifilm Wako) and 25 U/mL DNase I (Fujifilm Wako) through the bile duct (Figure S2C). The perfused pancreas was removed carefully from the intestine, stomach, spleen, and LNs, and dissociated for 30 min at 37℃. Then, 5 mL of DMEM containing 10% FBS and 1% PcSM was added to the dissociated pancreas and shaken for 10 seconds to completely dissociate the tissue. The digested pancreas was filtered through a 70-µm cell strainer with a syringe plunger. Red blood cell lysis was performed by incubation with ammonium-chloride-potassium lysis buffer for 4 min on ice. The digested pancreas was centrifuged and resuspended in 40% Percoll (GE Healthcare Life Science, Buckinghamshire, UK) layered on 70% Percoll, and centrifuged at 2200 rpm for 20 min. Cells were collected from the Percoll interface. Isolated cells were preincubated with CD16/32 blocker (clone 93; BioLegend) for 5 minutes at 4°C to prevent nonspecific staining, and stained for 20 min at 4℃ with antibodies. We added 1 mg/mL propidium iodide to the cell suspension to exclude dead cells (Figure S2C).

To stain IFN-γ and IL-17A, cells were stimulated with 25 ng/mL PMA (Fujifilm Wako) and 1 mg/mL ionomycin (Fujifilm Wako) for 6 hours at 37℃. Then, 10 mg/mL Brefeldin A (Fujifilm Wako) was added for the last 5 hours of incubation. Intracellular staining was performed using a Cytofix/Cytoperm Fixation/Permeabilization Solution kit (BD Biosciences, San Jose, CA, USA) according to the manufacturer’s instructions. The intracellular staining of Foxp3, Ki67, T-bet, Eomes, TCF-1, and Blimp-1, was performed using the Foxp3/Transcription factor staining buffer set (Invitrogen) according to the manufacturer’s instructions. Dead cells were excluded using a Zombie Aqua fixable viability kit (BioLegend). Stained cells were analyzed by FACSVerse flow cytometer (BD Biosciences) and data were analyzed using FlowJo software (Tree Star).

The following antibodies were used in this study. FITC anti-CD8a (53-6.7), FITC anti-CD62L (MEL-14), FITC anti-CD4 (RM4-5), FITC anti-RT1B (I-A g7) (OX-6), PE anti-CD4 (GK1.5), PE anti-CD3 (17A2), PE anti-Foxp3 (MF-14), PE anti-IFN-γ (XMG1.2), PE anti-Ki67 (16A8), PE anti-T-bet (4B10), PE anti-Blimp-1 (5E7), PE anti-CD40 (3/23), PerCP Cy5.5 anti-CD3 (17A2), PerCP Cy5.5 anti-CD44 (1M7), APC anti-CD19 (1D3), APC anti-CD44 (1M7), APC anti-CD25 (PC61), APC anti-Cxcr3 (CXCR3-173), APC anti-CD86 (GL-1), APC anti-CD103 (2E7), APC-Cy7 anti-CD3 (17A2), APC-Cy7 anti-TCR γ/δ (GL3), APC-Cy7 anti-CD45 (30-F11), APC-Cy7 anti-CD8a (53-6.7), PE-Cy7 anti-TCR γ/δ (GL3), PE-Cy7 anti-CD8a (53-6.7), PE-Cy7 anti-CD4 (RM4-5), PE-Cy7 anti-PD1 (RMP1-30), BV421 anti-H2-Kd (SF-1.1), BV421 anti-Cxcr3 (CXCR3-173), and BV510 anti-CD45 (30-F11) were purchased from BioLegend. PE anti-CD3 (500A2), PE anti-CD11b (M1/70), PerCP Cy5.5 anti-CD19 (1D3), PE anti-CD44 (1M7), PE-Cy7 anti-Eomes (Dan11mag), and PE-Cy7 anti-CD11c (N418) were purchased from eBioscience. BV450 anti-CD4 (RM4-5), APC anti-IL17A (TC11-18H10), and BV450 anti-CD11b (M1/70) were purchased from BD Biosciences. PE anti-TCF1 (C63D9) was purchased from Cell Signaling Technology.

### Quantitative PCR

mRNA was extracted using Isogen II (Nippon gene, Tokyo, Japan) and cDNA was synthesized using a High-capacity cDNA reverse transcription kit (Applied Biosystems, Foster City, CA, USA) according to the manufacturer’s instructions. PCR was performed on a PCR thermal cycler (Takara, Tokyo, Japan) and real-time PCR was performed using QuantStudio 3 (Thermo Fisher Scientific, Waltham, MA, USA). The relative quantification value is expressed as 2^−ΔΔ^Ct, where ΔΔ*Ct* is the difference between the mean *Ct* value of duplicate measurements of the sample and the endogenous *Actb* control. Acrb: primer set ID, MA050368 (Takara Bio); Cxcl10: primer set ID, MA118556 (Takara Bio); Cxcl9: forward, 5′-CCTAGTGATAAGGAATGCACGATG-3′, reverse, 5′-CTAGGCAGGTTTGATCTCCGTTC-3′; Cxcl13: forward, 5′-CATAGATCGGATTCAAGTTACGCC-3′, reverse,5′-GTAACCATTTGGCACGAGGATTC-3′; Fas: forward, 5′-CTGCGATTCTCCTGGCTGTGAA-3′, reverse, 5′-CAACAACCATAGGCGATTTCTGG-3′; and Isg15: forward, 5′-CATCCTGGTGAGGAACGAAAGG-3′, reverse, 5′-CTCAGCCAGAACTGGTCTTCGT-3′.

### RNA-sequencing analysis

To analyze CD44^hi^ IGRP CD8 T cells and CD8^+^ rDC in the PLN, NOD mice aged 6 weeks were used. Approximately 300-600 cells were sorted and used for the analysis. To analyze IGRP CD8 T cells in the pancreas, NOD mice aged 10-12 weeks were used. Approximately 600 cells were sorted and used for the analysis. Complementary DNA was prepared using a SMART-Seq v4 Ultra Low Input RNA Kit for Sequencing (Clontech Laboratories, CA, USA). Libraries were prepared using a Nextera XT DNA Library Prep Kit (Illumine, San Diego, CA, USA). Samples were run on a NovaSeq 6000 (Illumina). An average of 64 million paired reads were generated per sample. The accession number for the data is SRA: PRJNA992677. RNA-seq analysis was supported by Takara Bio Inc. (Japan).

### *Ex vivo* co-culture assay

To obtain FACS-sorted CD8^+^ rDCs, spleens, iLN, and PLN from non-diabetic 6w– 8w female mice were minced and enzymatically dissociated with 1 mg/mL collagenase D (Roche Applied Science) and 25 U/mL DNase I in DMEM for 30 min at 37°C, as described above. The digested tissues were filtered through a 70-µm cell strainer with a syringe plunger. Red blood cell lysis was performed by incubation with ammonium-chloride-potassium lysis buffer for 5 min on ice. The digested tissues were centrifuged and resuspended in 40% Percoll (GE Healthcare Life Science) layered on 70% Percoll, and centrifuged at 2200 rpm for 20 min. Cells were collected from the Percoll interface. Then, isolated cells were preincubated with CD16/32 blocker (93; BioLegend) for 5 minutes at 4°C to prevent nonspecific staining, and stained for 5 min at 4°C with biotinylated anti-CD3 antibody (17A2; BioLegend) to exclude CD3-positive T cells in the cell suspension using anti-biotin microbeads (Miltenyi Biotec) and LS columns (Miltenyi Biotec). Next, the cells were stained for 5 min at 4°C with biotinylated anti-CD19 antibody (1D3; BioLegend) to exclude CD19-positive B cells in the cell suspension using anti-biotin microbeads and LS columns. The cells were stained with antibodies (FITC anti-RT1B (OX-6), PE anti-CD11b (M1/70), APC anti-CD103 (2E7), APC-Cy7 anti-CD8a (53-6.7), PE-Cy7 anti-CD11c (N418), BV510 anti-CD45 (30-F11)) and sorted using MA900 (Sony). To obtain naïve 8.3 CD8 T cells, naïve CD8 T cells were isolated from the LNs of NY 8.3 NOD mice using two rounds of the naïve 8.3 CD8 T cell isolation kit (Miltenyi Biotec) with LS columns (Miltenyi Biotec), following the manufacturer’s instructions. Purified naïve 8.3 CD8 T cells were labeled using a CellTrace violet cell proliferation kit (Invitrogen) according to the manufacturer’s protocol. For the co-culture assay, 5×10^4^ sorted CD8^+^ rDC and 2.5×10^5^ CTV labeled naïve 8.3 CD8 T cells (DC:CD8 T ratio = 1:5) were mixed and cultured in U-bottom 96-well plates in RPMI-1640 with 10% FBS, 1% PcSM, 50 µM 2-mercaptoethanol, and 2µg IGRP_206-214_ (VYLKTNVFL). Cultures of CTV-labeled naïve 8.3 CD8 T cells without antigen-loaded DCs for 3d were used as a control. The cells were cultured at 37°C and 5% CO_2_ for 72 hours and then the dye dilution was analyzed using a FACSVerse flow cytometer (BD Biosciences), and data were analyzed using FlowJo software (Tree Star).

### *Ex vivo* cytotoxicity assay

To assess the effector functions of CD8 T cells against β-cells, *ex vivo* activated 8.3 CD8 T cells and NIT-1 cells were co-cultured as previously described (Zhang et al., 2014). Briefly, 8.3 CD8 T cells were isolated from the LNs of non-diabetic NY 8.3 NOD female mice using two rounds of the CD8 T cell isolation kit (Miltenyi Biotec) with LS columns (Miltenyi Biotec), following the manufacturer’s instructions. The isolated cells were cultured in U-bottom 96-well plates in RPMI-1640 with 10% FBS, 1% PcSM, 50 µM 2-mercaptoethanol, 30 U/mL IL-2, 5 ng/mL IL-12, and Dynabeads Mouse T-activator CD3/CD28 (Thermo Fisher) (8.3 CD8 T:beads = 1:1) in the presence or absence of BMS-986165 at 37°C and 5% CO_2_ for 72 hours. Then, 1×10^5^ NIT-1 cells were cultured in flat-bottom 96-well plates in F12K with 10% FBS, 1% PcSM, and incubated at 37°C and 5% CO_2_ overnight. For the co-culture cytotoxicity assay, *ex vivo* activated 8.3 CD8 T cells were added to a plate with NIT-1 cells, and co-cultured in F12K with 10% FBS, 1% PcSM, and 2 µg IGRP_206-214_ (VYLKTNVFL) at 37°C and 5% CO_2_ for 24 hours (NIT-1:CD8 T ratio = 1:0.1, 1:1, 1:2). After incubation, the supernatants were removed and washed three times with PBS(-) to remove 8.3 CD8 T cells. After washing, 100 µL of F12K with 10% FBS, 1% PcSM, and 20 µL of CellTiter 96 (Promega, Tokyo, Japan) was added and incubated at 37°C and 5% CO_2_ for 2 hours. The absorbance was determined at 450 nm. The % killing was calculated as follows: % killing = 100 − ((Abs of NIT-1 cells cultured with 8.3 CD8+ T cells)/(Abs of untreated NIT-1 cells) × 100).

### BMS-986165 treatment

BMS-986165 (deucravacitinib) (6-cyclopropaneamido-4-{[2-methoxy-3-(1-methyl-1H-1,2,4-triazol-3-yl)phenyl]amino}-N-(2H3)methylpyridazine-3-carboxamide) was purchased from Selleck Biotech (Tokyo, Japan). To test the inhibitory effects of BMS-986165 on the development of autoimmune T1D, wild type NOD mice were dosed by oral gavage once daily with BMS-986165 30 mg/kg in vehicle (EtOH:TPGS:PEG300, 5:5:90) or vehicle alone as previously described (Wrobleski et al., 2019). Blood glucose levels were measured weekly in tail vein blood using Glutest Ai (Sanwa).

### Data analysis

Heatmaps were generated with the R (3.6.0) package pheatmap, and clustering was performed using the word.D2 method. Enrichment analysis was performed by GeneTrail (Gerstner et al., 2020). PCA analysis was performed using ClustVis (Metsalu and Vilo, 2015) and RNAseqChef (Etoh and Nakao, 2023). Volcano plots were generated using ggVolcanoR (Mullan et al., 2021). Likelihood ratio tests were performed to determine differentially expressed genes between samples using RNAseqChef

### Statistical analysis

Data are expressed as the means ± s.e.m. Statistical significance between two groups was determined by unpaired Student’s *t*-test. For the analysis of diabetes incidence, Kaplan-Meier survival curves were estimated by the log rank test. Insulitis levels were analyzed by the Mann-Whitney *U*-test. *p* < 0.05 was considered statistically significant. Statistical analysis was performed using Prism 9 (GraphPad Software). Differentially expressed genes between pairwise comparisons were performed using DESeq2 (FDR < 0.1, fold change > 1.5) in iDEP.96 (Ge et al., 2018). For the cluster analysis of IGRP CD8 T cells in the PLN and pancreas, likelihood ratio tests were used to determine differentially expressed genes between the samples (FDR < 0.1, fold change > 1.5) using RNAseqChef (Etoh and Nakao, 2023). We did not use *a priori* statistical methods to determine the sample size. Sample size sufficiency was based on previous experiments from our laboratory and others. We were not blinded to the allocation of mice, samples, and outcome data analyses.

## Author contributions

Conceptualization, K.M., K.A. and S.N.; Investigation, K.M.; writing - original draft preparation, K.M.; writing - review and editing, H.T., H.M., S.A., N.A., H.K., K.A. and S.N.; supervision, S.N.; project administration, K.M.; resources, S.A., N.A., and H.K.; funding acquisition, K.M., and S.N. All authors have read and agreed to the published version of the manuscript.

## Institutional Review Board Statement

Not applicable.

## Informed Consent Statement

Not applicable.

## Data Availability Statement

GSE235670, PRJNA992677

## Acknowledgments

We sincerely thank Shinichiro Sawa, Kota Yanagitani, Shin-ichi Koizumi, Naoto Noguchi, and Shinya Hatano for their counsel on experiments, and Yumiko Kitada, Akiko Takano, Chieko Ogawa, Saori Fuchigami, Rasheda Perveen, and Yoshifumi Morita for helping to prepare the manuscript. We appreciate technical support from the Analytical Research Center for Experimental Sciences, Saga University. We thank Edanz (https://jp.edanz.com/ac) for editing a draft of this manuscript.

## Abbreviations

DC: dendritic cell
NOD: non-obese diabetic
TMRE: tetramethylrhodamine ethyl ester perchlorate Tyk2, tyrosine kinase 2

## REFERENCES

Aizu, K., W. Li, T. Yajima, T. Arai, K. Shimoda, Y. Nimura, and Y. Yoshikai. 2006. An important role of Tyk2 in APC function of dendritic cells for priming CD8+ T cells producing IFN-γ. Eur. J. Immunol. 36:3060–3070. doi:10.1002/eji.200636173.

Bradfield, J.P., H.Q. Qu, K. Wang, H. Zhang, P.M. Sleiman, C.E. Kim, F.D. Mentch, H. Qiu, J.T. Glessner, K.A. Thomas, E.C. Frackelton, R.M. Chiavacci, M. Imielinski, D.S. Monos, R. Pandey, M. Bakay, S.F.A. Grant, C. Polychronakos, and H. Hakonarson. 2011. A genome-wide meta-analysis of six type 1 diabetes cohorts identifies multiple associated loci. PLoS Genet. 7:1002293. doi:10.1371/journal.pgen.1002293.

Burke, J.R., L. Cheng, K.M. Gillooly, J. Strnad, A. Zupa-Fernandez, I.M. Catlett, Y. Zhang, E.M. Heimrich, K.W. McIntyre, M.D. Cunningham, J.A. Carman, X. Zhou, D. Banas, C. Chaudhry, S. Li, C. D’Arienzo, A. Chimalakonda, X.X. Yang, J.H. Xie, J. Pang, Q. Zhao, S.M. Rose, J. Huang, R.M. Moslin, S.T. Wrobleski, D.S. Weinstein, and L.M. Salter-Cid. 2019. Autoimmune pathways in mice and humans are blocked by pharmacological stabilization of the TYK2 pseudokinase domain. Sci. Transl. Med. 11:1736. doi:10.1126/scitranslmed.aaw1736.

Chiou, J., R.J. Geusz, M.L. Okino, J.Y. Han, M. Miller, R. Melton, E. Beebe, P. Benaglio, S. Huang, K. Korgaonkar, S. Heller, A. Kleger, S. Preissl, D.U. Gorkin, M. Sander, and K.J. Gaulton. 2021. Interpreting type 1 diabetes risk with genetics and single-cell epigenomics. Nature. 594:398–402. doi:10.1038/s41586-021-03552-w.

Collins, C., F.W.L. Tsui, and M.J. Shulman. 2002. Induction of the chemokine receptor CXCR3 on TCR-stimulated T cells: Dependence on the release from persistent TCR-triggering and requirement for IFN-γ stimulation. Eur. J. Immunol. 32:1792– 1801. doi:10.1002/1521-4141(200206):6<1792::AID-IMMU1792>3.0.CO;2-0.

Coppieters, K.T., T. Boettler, and M. von Herrath. 2012. Virus Infections in Type 1 Diabetes. Cold Spring Harb. Perspect. Med. 2:a007682–a007682. doi:10.1101/cshperspect.a007682.

Cullen, S.P., C.M. Henry, C.J. Kearney, S.E. Logue, M. Feoktistova, G.A. Tynan, E.C. Lavelle, M. Leverkus, and S.J. Martin. 2013. Fas/CD95-Induced Chemokines Can Serve as “ Find-Me” Signals for Apoptotic Cells. Mol. Cell. 49:1034–1048. doi:10.1016/j.molcel.2013.01.025.

Derecka, M., A. Gornicka, S.B. Koralov, K. Szczepanek, M. Morgan, V. Raje, J. Sisler, Q. Zhang, D. Otero, J. Cichy, K. Rajewsky, K. Shimoda, V. Poli, B. Strobl, S. Pellegrini, T.E. Harris, P. Seale, A.P. Russell, A.J. McAinch, P.E. O’Brien, S.R. Keller, C.M. Croniger, T. Kordula, and A.C. Larner. 2012. Tyk2 and stat3 regulate brown adipose tissue differentiation and obesity. Cell Metab. 16:814–824. doi:10.1016/j.cmet.2012.11.005.

DiMeglio, L.A., C. Evans-Molina, and R.A. Oram. 2018. Type 1 diabetes. Lancet. 391:2449–2462. doi:10.1016/S0140-6736(18)31320-5.

Ellis, J.A., A.S. Kemp, and A.L. Ponsonby. 2014. Gene-environment interaction in autoimmune disease. Expert Rev. Mol. Med. 16:1–23. doi:10.1017/erm.2014.5.

Etoh, K., and M. Nakao. 2023. A web-based integrative transcriptome analysis, RNAseqChef, uncovers the cell/tissue type-dependent action of sulforaphane. J. Biol. Chem. 299:104810. doi:10.1016/j.jbc.2023.104810.

Flodström, M., A. Maday, D. Balakrishna, M.M. Cleary, A. Yoshimura, and N. Sarvetnick. 2002. Target cell defense prevents the development of diabetes after viral infection. Nat. Immunol. 3:373–382. doi:10.1038/ni771.

Fukui, T., T. Kobayashi, E. Jimbo, K. Aida, A. Shimada, Y. Oikawa, Y. Mori, T. Fujii, R. Koyama, K. Kobayashi, A. Takeshita, and S. Yagihashi. 2023. Bi-glandular and persistent enterovirus infection and distinct changes of the pancreas in slowly progressive type 1 diabetes mellitus. Sci. Rep. 13:1–16. doi:10.1038/s41598-023-33011-7.

Gagnerault, M.C., J.J. Luan, C. Lotton, and F. Lepault. 2002. Pancreatic lymph nodes are required for priming of β cell reactive T cells in NOD mice. J. Exp. Med. 196:369–377. doi:10.1084/jem.20011353.

Ge, S.X., E.W. Son, and R. Yao. 2018. iDEP: An integrated web application for differential expression and pathway analysis of RNA-Seq data. BMC Bioinformatics. 19:1–24. doi:10.1186/s12859-018-2486-6.

Gearty, S. V., F. Dündar, P. Zumbo, G. Espinosa-Carrasco, M. Shakiba, F.J. Sanchez-Rivera, N.D. Socci, P. Trivedi, S.W. Lowe, P. Lauer, N. Mohibullah, A. Viale, T.P. DiLorenzo, D. Betel, and A. Schietinger. 2022. An autoimmune stem-like CD8 T cell population drives type 1 diabetes. Nature. 602:156–161. doi:10.1038/s41586-021-04248-x.

Gerstner, N., T. Kehl, K. Lenhof, A. Müller, C. Mayer, L. Eckhart, N.L. Grammes, C. Diener, M. Hart, O. Hahn, J. Walter, T. Wyss-Coray, E. Meese, A. Keller, and H.P. Lenhof. 2020. GeneTrail 3: Advanced high-throughput enrichment analysis. Nucleic Acids Res. 48:W515–W520. doi:10.1093/NAR/GKAA306.

Girdhar, K., Q. Huang, I.T. Chow, T. Vatanen, C. Brady, A. Raisingani, P. Autissier, M.A. Atkinson, W.W. Kwok, C. Ronald Kahn, and E. Altindis. 2022. A gut microbial peptide and molecular mimicry in the pathogenesis of type 1 diabetes. Proc. Natl. Acad. Sci. U. S. A. 119:1–12. doi:10.1073/pnas.2120028119.

Gorman, J.A., C. Hundhausen, J.S. Errett, A.E. Stone, E.J. Allenspach, Y. Ge, T. Arkatkar, C. Clough, X. Dai, S. Khim, K. Pestal, D. Liggitt, K. Cerosaletti, D.B. Stetson, R.G. James, M. Oukka, P. Concannon, M. Gale, J.H. Buckner, and D.J. Rawlings. 2017. The A946T variant of the RNA sensor IFIH1 mediates an interferon program that limits viral infection but increases the risk for autoimmunity. Nat. Immunol. 18:744–752. doi:10.1038/ni.3766.

Gracey, E., D. Hromadová, M. Lim, Z. Qaiyum, M. Zeng, Y. Yao, A. Srinath, Y. Baglaenko, N. Yeremenko, W. Westlin, C. Masse, M. Müller, B. Strobl, W. Miao, and R.D. Inman. 2020. TYK2 inhibition reduces type 3 immunity and modifies disease progression in murine spondyloarthritis. J. Clin. Invest. 130:1863–1878. doi:10.1172/JCI126567.

Greb, J.E., A.M. Goldminz, J.T. Elder, M.G. Lebwohl, D.D. Gladman, J.J. Wu, N.N. Mehta, A.Y. Finlay, and A.B. Gottlieb. 2016. Psoriasis. Nat. Rev. Dis. Prim. 2. doi:10.1038/nrdp.2016.82.

Gregory, G.A., T.I.G. Robinson, S.E. Linklater, F. Wang, S. Colagiuri, C. de Beaufort, K.C. Donaghue, D.J. Magliano, J. Maniam, T.J. Orchard, P. Rai, and G.D. Ogle. 2022. Global incidence, prevalence, and mortality of type 1 diabetes in 2021 with projection to 2040: a modelling study. *lancet*. Diabetes Endocrinol. 10:741–760. doi:10.1016/S2213-8587(22)00218-2.

Hamaguchi, K., H.R. Gaskins, and E.H. Leiter. 1991. NIT-1, a pancreatic β-cell line established from a transgenic NOD/Lt mouse. Diabetes. 40:842–849. doi:10.2337/diab.40.7.842.

Hashiguchi, T., A. Oyamada, K. Sakuraba, K. Shimoda, K.I. Nakayama, Y. Iwamoto, Y. Yoshikai, and H. Yamada. 2014. Tyk2-dependent bystander activation of conventional and nonconventional Th1 cell subsets contributes to innate host defense against Listeria monocytogenes infection. J. Immunol. 192:4739–47. doi:10.4049/jimmunol.1303067.

Herold, K.C., B.N. Bundy, S.A. Long, J.A. Bluestone, L.A. DiMeglio, M.J. Dufort, S.E. Gitelman, P.A. Gottlieb, J.P. Krischer, P.S. Linsley, J.B. Marks, W. Moore, A. Moran, H. Rodriguez, W.E. Russell, D. Schatz, J.S. Skyler, E. Tsalikian, D.K. Wherrett, A.-G. Ziegler, and C.J. Greenbaum. 2019. An Anti-CD3 Antibody, Teplizumab, in Relatives at Risk for Type 1 Diabetes. N. Engl. J. Med. 381:603–613. doi:10.1056/nejmoa1902226.

Hildner, K., B.T. Edelson, W.E. Purtha, M. Diamond, H. Matsushita, M. Kohyama, B. Calderon, B.U. Schraml, E.R. Unanue, M.S. Diamond, R.D. Schreiber, T.L. Murphy, and K.M. Murphy. 2008. Batf3 deficiency reveals a critical role for CD8α+ dendritic cells in cytotoxic T cell immunity. Science (80-. ). 322:1097–1100. doi:10.1126/science.1164206.

Horwitz, D.A., T.M. Fahmy, C.A. Piccirillo, and A. La Cava. 2019. Rebalancing Immune Homeostasis to Treat Autoimmune Diseases. Trends Immunol. 40:888– 908. doi:10.1016/j.it.2019.08.003.

Hosokawa, Y., T. Hanafusa, and A. Imagawa. 2019. Pathogenesis of fulminant type 1 diabetes: Genes, viruses and the immune mechanism, and usefulness of patient-derived induced pluripotent stem cells for future research. J. Diabetes Investig. 10:1158–1164. doi:10.1111/jdi.13091.

Imagawa, A., T. Hanafusa, J.I. Miyagawa, and Y. Matsuzawa. 2000. A proposal of three distinct subtypes of type 1 diabetes mellitus based on clinical and pathological evidence. Ann. Med. 32:539–543. doi:10.3109/07853890008998833.

Imagawa, A., T. Hanafusa, Y. Uchigata, A. Kanatsuka, E. Kawasaki, T. Kobayashi, A. Shimada, I. Shimizu, T. Toyoda, T. Maruyama, and H. Makino. 2003. Fulminant type 1 diabetes: A nationwide survey in Japan. Diabetes Care. 26:2345–2352. doi:10.2337/diacare.26.8.2345.

Ishizaki, M., T. Akimoto, R. Muromoto, M. Yokoyama, Y. Ohshiro, Y. Sekine, H. Maeda, K. Shimoda, K. Oritani, and T. Matsuda. 2011. Involvement of Tyrosine Kinase-2 in Both the IL-12/Th1 and IL-23/Th17 Axes In Vivo. J. Immunol. 187:181–189. doi:10.4049/jimmunol.1003244.

Ishizaki, M., R. Muromoto, T. Akimoto, Y. Sekine, S. Kon, M. Diwan, H. Maeda, S. Togi, K. Shimoda, K. Oritani, and T. Matsuda. 2014. Tyk2 is a therapeutic target for psoriasis-like skin inflammation. Int. Immunol. 26:257–267. doi:10.1093/intimm/dxt062.

Izumi, K., K. Mine, Y. Inoue, M. Teshima, S. Ogawa, Y. Kai, T. Kurafuji, K. Hirakawa, D. Miyakawa, H. Ikeda, A. Inada, M. Hara, H. Yamada, K. Akashi, Y. Niho, K. Ina, T. Kobayashi, Y. Yoshikai, K. Anzai, T. Yamashita, H. Minagawa, S. Fujimoto, H. Kurisaki, K. Shimoda, H. Katsuta, and S. Nagafuchi. 2015. Reduced Tyk2 gene expression in β-cells due to natural mutation determines susceptibility to virus-induced diabetes. Nat. Commun. 6:6748. doi:10.1038/ncomms7748.

Jayaraman, S. 2011. Assessment of beta cell viability. Curr. Protoc. Cytom. doi:10.1002/0471142956.cy0627s55.

Jenson, A.B., H. Rosenberg, and A.L. Notkins. 1980. Pancreatic islet-cell damage in children with fatal viral infections. Lancet. 316:354–358.

Kim, H., J. Perovanovic, A. Shakya, Z. Shen, C.N. German, A. Ibarra, J.L. Jafek, N.P. Lin, B.D. Evavold, D.H.C. Chou, P.E. Jensen, X. He, and D. Tantin. 2021. Targeting transcriptional coregulator OCA-B/Pou2af1 blocks activated autoreactive T cells in the pancreas and type 1 diabetes. J. Exp. Med. 218. doi:10.1084/jem.20200533.

Knip, M., and O. Simell. 2012. Environmental triggers of type 1 diabetes. Cold Spring Harb Perspect Med. 2:a007690. doi:10.1101/cshperspect.a007690.

Kreins, A.Y., M.J. Ciancanelli, S. Okada, X.-F. Kong, N. Ramírez-Alejo, S.S. Kilic, J. El Baghdadi, S. Nonoyama, S.A. Mahdaviani, F. Ailal, A. Bousfiha, D. Mansouri, E. Nievas, C.S. Ma, G. Rao, A. Bernasconi, H. Sun Kuehn, J. Niemela, J. Stoddard, P. Deveau, A. Cobat, S. El Azbaoui, A. Sabri, C.K. Lim, M. Sundin, D.T. Avery, R. Halwani, A. V Grant, B. Boisson, D. Bogunovic, Y. Itan, M. Moncada-Velez, R. Martinez-Barricarte, M. Migaud, C. Deswarte, L. Alsina, D. Kotlarz, C. Klein, I. Muller-Fleckenstein, B. Fleckenstein, V. Cormier-Daire, S. Rose-John, C. Picard, L. Hammarstrom, A. Puel, S. Al-Muhsen, L. Abel, D. Chaussabel, S.D. Rosenzweig, Y. Minegishi, S.G. Tangye, J. Bustamante, J.-L. Casanova, and S. Boisson-Dupuis. 2015. Human TYK2 deficiency: Mycobacterial and viral infections without hyper-IgE syndrome. J. Exp. Med. 212:1641–62. doi:10.1084/jem.20140280.

Kuriya, G., T. Uchida, S. Akazawa, M. Kobayashi, K. Nakamura, T. Satoh, I. Horie, E. Kawasaki, H. Yamasaki, L. Yu, Y. Iwakura, H. Sasaki, Y. Nagayama, A. Kawakami, and N. Abiru. 2013. Double deficiency in IL-17 and IFN-γ signalling significantly suppresses the development of diabetes in the NOD mouse. Diabetologia. 56:1773–1780. doi:10.1007/s00125-013-2935-8.

Kurts, C., B.W.S. Robinson, and P.A. Knolle. 2010. Cross-priming in health and disease. Nat. Rev. Immunol. 10:403–414. doi:10.1038/nri2780.

Kuznetsov, A. V., R. Margreiter, A. Amberger, V. Saks, and M. Grimm. 2011. Changes in mitochondrial redox state, membrane potential and calcium precede mitochondrial dysfunction in doxorubicin-induced cell death. Biochim. Biophys. Acta - Mol. Cell Res. 1813:1144–1152. doi:10.1016/j.bbamcr.2011.03.002.

Lalwani, A., J. Warren, D. Liuwantara, W.J. Hawthorne, P.J. O’Connell, F.J. Gonzalez, R.A. Stokes, J. Chen, D.R. Laybutt, M.E. Craig, M.M. Swarbrick, C. King, and J.E. Gunton. 2019. β Cell Hypoxia-Inducible Factor-1α Is Required for the Prevention of Type 1 Diabetes. Cell Rep. 27:2370–2384.e6. doi:10.1016/j.celrep.2019.04.086.

Lazarevic, V., L.H. Glimcher, and G.M. Lord. 2013. T-bet: A bridge between innate and adaptive immunity. Nat. Rev. Immunol. 13:777–789. doi:10.1038/nri3536.

Leitner, N.R., A. Witalisz-Siepracka, B. Strobl, and M. Müller. 2017. Tyrosine kinase 2 – Surveillant of tumours and bona fide oncogene. Cytokine. 89:209–218. doi:10.1016/j.cyto.2015.10.015.

Li, W., H. Yamada, T. Yajima, R. Nakagawa, K. Shimoda, K. Nakayama, and Y. Yoshikai. 2007. Tyk2 Signaling in Host Environment Plays an Important Role in Contraction of Antigen-Specific CD8+ T Cells following a Microbial Infection. J. Immunol. 178:4482–4488. doi:10.4049/jimmunol.178.7.4482.

Luo, X., K.C. Herold, and S.D. Miller. 2010. Immunotherapy of Type 1 Diabetes: Where Are We and Where Should We Be Going? Immunity. 32:488–499. doi:10.1016/j.immuni.2010.04.002.

Maruyama, C., H. Suemizu, S. Tamamushi, S. Kimoto, N. Tamaoki, and Y. Ohnishi. 2002. Genotyping the mouse severe combined immunodeficiency mutation using the polymerase chain reaction with confrontingtwo-pair primers (PCR-CTPP). Exp. Anim. 51:391–393. doi:10.1538/expanim.51.391.

Metsalu, T., and J. Vilo. 2015. ClustVis: A web tool for visualizing clustering of multivariate data using Principal Component Analysis and heatmap. Nucleic Acids Res. 43:W566–W570. doi:10.1093/nar/gkv468.

Mine, K., K. Hirakawa, S. Kondo, M. Minami, A. Okada, N. Tsutsu, Y. Yokogawa, Y. Hibio, F. Kojima, S. Fujimoto, H. Kurisaki, K. Anzai, Y. Yoshikai, and S. Nagafuchi. 2017. Subtyping of Type 1 Diabetes as Classified by Anti-GAD Antibody, IgE Levels, and Tyrosine kinase 2 (TYK2) Promoter Variant in the Japanese. EBioMedicine. 23:46–51. doi:10.1016/j.ebiom.2017.08.012.

Mine, K., S. Nagafuchi, S. Hatano, K. Tanaka, H. Mori, H. Takahashi, K. Anzai, and Y. Yoshikai. 2020a. Impaired upregulation of Stat2 gene restrictive to pancreatic β-cells is responsible for virus-induced diabetes in DBA/2 mice. Biochem. Biophys. Res. Commun. 521:853–860. doi:10.1016/j.bbrc.2019.10.193.

Mine, K., Y. Yoshikai, H. Takahashi, H. Mori, K. Anzai, and S. Nagafuchi. 2020b. Genetic susceptibility of the host in virus-induced diabetes. Microorganisms. 8:1–20. doi:10.3390/microorganisms8081133.

Mori, H., H. Takahashi, K. Mine, K. Higashimoto, K. Inoue, M. Kojima, S. Kuroki, T. Eguchi, Y. Ono, S. Inuzuka, H. Soejima, S. Nagafuchi, and K. Anzai. 2021. TYK2 promoter variant is associated with impaired insulin secretion and lower insulin resistance in japanese type 2 diabetes patients. Genes (Basel). 12. doi:10.3390/genes12030400.

Moriwaki, M., N. Itoh, J. Miyagawa, K. Yamamoto, A. Imagawa, K. Yamagata, H. Iwahashi, H. Nakajima, M. Namba, S. Nagata, T. Hanafusa, and Y. Matsuzawa. 1999. Fas and Fas ligand expression in inflamed islets in pancreas sections of patients with recent-onset Type I diabetes mellitus. Diabetologia. 42:1332–1340. doi:10.1007/s001250051446.

Mullan, K.A., L.M. Bramberger, P.R. Munday, G. Goncalves, J. Revote, N.A. Mifsud, P.T. Illing, A. Anderson, P. Kwan, A.W. Purcell, and C. Li. 2021. ggVolcanoR: A Shiny app for customizable visualization of differential expression datasets. Comput. Struct. Biotechnol. J. 19:5735–5740. doi:10.1016/j.csbj.2021.10.020.

Müller, J.A., R. Groß, C. Conzelmann, J. Krüger, U. Merle, J. Steinhart, T. Weil, L. Koepke, C.P. Bozzo, C. Read, G. Fois, T. Eiseler, J. Gehrmann, J. van Vuuren, I.M. Wessbecher, M. Frick, I.G. Costa, M. Breunig, B. Grüner, L. Peters, M. Schuster, S. Liebau, T. Seufferlein, S. Stenger, A. Stenzinger, P.E. MacDonald, F. Kirchhoff, K.M.J. Sparrer, P. Walther, H. Lickert, T.F.E. Barth, M. Wagner, J. Münch, S. Heller, and A. Kleger. 2021. SARS-CoV-2 infects and replicates in cells of the human endocrine and exocrine pancreas. Nat. Metab. 3:149–165. doi:10.1038/s42255-021-00347-1.

Muromoto, R., K. Shimoda, K. Oritani, and T. Matsuda. 2021. Therapeutic advantage of tyk2 inhibition for treating autoimmune and chronic inflammatory diseases. Biol. Pharm. Bull. 44:1585–1592. doi:10.1248/bpb.b21-00609.

Nagafuchi, S., Y. Kamada-Hibio, K. Hirakawa, N. Tsutsu, M. Minami, A. Okada, K. Kai, M. Teshima, A. Moroishi, Y. Murakami, Y. Umeno, Y. Yokogawa, K. Kogawa, K. Izumi, K. Anzai, R. Iwakiri, K. Hamaguchi, N. Sasaki, S. Nohara, E. Yoshida, M. Harada, K. Akashi, T. Yanase, J. Ono, T. Okeda, R. Fujimoto, K. Ihara, T. Hara, Y. Kikuchi, M. Iwase, T. Kitazono, F. Kojima, S. Kono, H. Kurisaki, S. Kondo, and H. Katsuta. 2015. TYK2 Promoter Variant and Diabetes Mellitus in the Japanese. EBioMedicine. 2:744–749. doi:10.1016/j.ebiom.2015.05.004.

O’Shea, J.J., D.M. Schwartz, A. V. Villarino, M. Gadina, I.B. McInnes, and A. Laurence. 2015. The JAK-STAT pathway: Impact on human disease and therapeutic intervention. Annu. Rev. Med. 66:311–328. doi:10.1146/annurev-med-051113-024537.

Onengut-Gumuscu, S., W.M. Chen, O. Burren, N.J. Cooper, A.R. Quinlan, J.C. Mychaleckyj, E. Farber, J.K. Bonnie, M. Szpak, E. Schofield, P. Achuthan, H. Guo, M.D. Fortune, H. Stevens, N.M. Walker, L.D. Ward, A. Kundaje, M. Kellis, M.J. Daly, J.C. Barrett, J.D. Cooper, P. Deloukas, J.A. Todd, C. Wallace, P. Concannon, and S.S. Rich. 2015. Fine mapping of type 1 diabetes susceptibility loci and evidence for colocalization of causal variants with lymphoid gene enhancers. Nat. Genet. 47:381–386. doi:10.1038/ng.3245.

Oyamada, A., H. Ikebe, M. Itsumi, H. Saiwai, S. Okada, K. Shimoda, Y. Iwakura, K.I. Nakayama, Y. Iwamoto, Y. Yoshikai, and H. Yamada. 2009. Tyrosine Kinase 2 Plays Critical Roles in the Pathogenic CD4 T Cell Responses for the Development of Experimental Autoimmune Encephalomyelitis. J. Immunol. 183:7539–7546. doi:10.4049/jimmunol.0902740.

Papp, K., K. Gordon, D. Thaçi, A. Morita, M. Gooderham, P. Foley, I.G. Girgis, S. Kundu, and S. Banerjee. 2018. Phase 2 Trial of Selective Tyrosine Kinase 2 Inhibition in Psoriasis. N. Engl. J. Med. 379:1313–1321. doi:10.1056/nejmoa1806382.

Reed, J.C., and K.C. Herold. 2015. Thinking bedside at the bench: The NOD mouse model of T1DM. Nat. Rev. Endocrinol. 11:308–314. doi:10.1038/nrendo.2014.236.

Shimoda, K., K. Kato, K. Aoki, T. Matsuda, A. Miyamoto, M. Shibamori, M. Yamashita, A. Numata, K. Takase, S. Kobayashi, S. Shibata, Y. Asano, H. Gondo, K. Sekiguchi, K. Nakayama, T. Nakayama, T. Okamura, S. Okamura, Y. Niho, and K. Nakayama. 2000a. Tyk2 plays a restricted role in IFN alpha signaling, although it is required for IL-12-mediated T cell function. Immunity. 13:561–571. doi:10.1016/S1074-7613(00)00055-8.

Shimoda, K., K. Kato, K. Aoki, T. Matsuda, A. Miyamoto, M. Shibamori, M. Yamashita, A. Numata, K. Takase, S. Kobayashi, S. Shibata, Y. Asano, H. Gondo, K. Sekiguchi, K. Nakayama, T. Nakayama, T. Okamura, S. Okamura, Y. Niho, and K. ichi Nakayama. 2000b. Tyk2 plays a restricted role in IFNα signaling, although it is required for IL-12-mediated T cell function. Immunity. 13:561–571. doi:10.1016/S1074-7613(00)00055-8.

Starbeck-Miller, G.R., H.H. Xue, and J.T. Harty. 2014. IL-12 and type I interferon prolong the division of activated CD8 T cells by maintaining high-affinity IL-2 signaling in vivo. J. Exp. Med. 211:105–120. doi:10.1084/jem.20130901.

Stone, V.M., M.M. Hankaniemi, O.H. Laitinen, A.B. Sioofy-Khojine, A. Lin, I.M. Diaz Lozano, M.A. Mazur, V. Marjomäki, K. Loré, H. Hyöty, V.P. Hytönen, and M. Flodström-Tullberg. 2020. A hexavalent Coxsackievirus B vaccine is highly immunogenic and has a strong protective capacity in mice and nonhuman primates.

Strober, B., D. Thaçi, H. Sofen, L. Kircik, K.B. Gordon, P. Foley, P. Rich, C. Paul, J. Bagel, E. Colston, J. Throup, S. Kundu, C. Sekaran, M. Linaberry, S. Banerjee, and K.A. Papp. 2023. Deucravacitinib versus placebo and apremilast in moderate to severe plaque psoriasis: Efficacy and safety results from the 52-week, randomized, double-blinded, phase 3 Program fOr Evaluation of TYK2 inhibitor psoriasis second trial. J. Am. Acad. Dermatol. 88:40–51. doi:10.1016/j.jaad.2022.08.061.

Uno, S., A. Imagawa, K. Saisho, K. Okita, H. Iwahashi, T. Hanafusa, and I. Shimomura. 2010. Expression of chemokines, CXC chemokine ligand 10 (CXCL10) and CXCR3 in the inflamed islets of patients with recent-onset autoimmune type 1 diabetes. Endocr. J. 57:991–6. doi:JST.JSTAGE/endocrj/K10E-076 [pii].

Verdaguer, J., D. Schmidt, A. Amrani, B. Anderson, N. Averill, and P. Santamaria. 1997. Spontaneous Autoimmune Diabetes in Monoclonal T Cell Nonobese Diabetic Mice. 186. 1663–1676 pp.

Vreugdenhil, G.R., N.C. Schloot, A. Hoorens, C. Rongen, D.G. Pipeleers, W.J.G. Melchers, B.O. Roep, and J.M.D. Galama. 2000. Acute onset of type I diabetes mellitus after severe echovirus 9 infection: Putative pathogenic pathways. Clin. Infect. Dis. 31:1025–1031. doi:10.1086/318159.

Works, M.G., F. Yin, C.C. Yin, Y. Yiu, K. Shew, T.-T. Tran, N. Dunlap, J. Lam, T. Mitchell, J. Reader, P.L. Stein, and A. D’Andrea. 2014. Inhibition of TYK2 and JAK1 Ameliorates Imiquimod-Induced Psoriasis-like Dermatitis by Inhibiting IL-22 and the IL-23/IL-17 Axis. J. Immunol. 193:3278–3287. doi:10.4049/jimmunol.1400205.

Wrobleski, S.T., R. Moslin, S. Lin, Y. Zhang, S. Spergel, J. Kempson, J.S. Tokarski, J. Strnad, A. Zupa-Fernandez, L. Cheng, D. Shuster, K. Gillooly, X. Yang, E. Heimrich, K.W. McIntyre, C. Chaudhry, J. Khan, M. Ruzanov, J. Tredup, D. Mulligan, D. Xie, H. Sun, C. Huang, C. D’Arienzo, N. Aranibar, M. Chiney, A. Chimalakonda, W.J. Pitts, L. Lombardo, P.H. Carter, J.R. Burke, and D.S. Weinstein. 2019. Highly Selective Inhibition of Tyrosine Kinase 2 (TYK2) for the Treatment of Autoimmune Diseases: Discovery of the Allosteric Inhibitor BMS-986165. J. Med. Chem. 62:8973–8995. doi:10.1021/acs.jmedchem.9b00444.

Yuan, S., L. Wang, H. Zhang, F. Xu, X. Zhou, L. Yu, J. Sun, J. Chen, H. Ying, X. Xu, Y. Yu, A. Spiliopoulou, X. Shen, J. Wilson, D. Gill, E. Theodoratou, S.C. Larsson, and X. Li. 2023. Mendelian randomization and clinical trial evidence supports TYK2 inhibition as a therapeutic target for autoimmune diseases. eBioMedicine. 89:104488. doi:10.1016/j.ebiom.2023.104488.

Zhang, L., J.N. Manirarora, and C.H. Wei. 2014. Evaluation of immunosuppressive function of regulatory T cells using a novel in vitro cytotoxicity assay. Cell Biosci. 4. doi:10.1186/2045-3701-4-51.

Zmuda, E.J., C.A. Powell, and T. Hai. 2011. A method for murine islet isolation and subcapsular kidney transplantation. J. Vis. Exp. 1–11. doi:10.3791/2096.

