## Supplemental figures for "Tyk2-mediated signaling promotes the development of autoreactive CD8^+^ CTLs and autoimmune type 1 diabetes"

### SUPPLEMENTARY FIGURE LEGENDS

**Figure S1. Establishing *Tyk2*KO.NOD mice and the analysis of purified β-cells.**
(A) PCR analysis of IDDM loci in *Tyk2*KO.NOD mice. (B) Body weight gain in *Tyk2*$^{+/+}$ (n=14), *Tyk2*$^{+/-}$ (n=10), and *Tyk2*$^{-/-}$ NOD female mice (n=13). (C) Representative hematoxylin and eosin staining of islets with or without insulitis. Score 0, no insulitis; Score 1, peri-insulitis; Score 2, infiltrative insulitis less than 50% of the islet area; and Score 3, infiltrative insulitis more than 50% of the islet area. (D) Flow cytometry plots and graphs show the frequency of naïve (CD44$^{lo}$ CD62L$^{+}$), effector memory (CD44$^{hi}$ CD62L$^{-}$), and central memory (CD44$^{hi}$ CD62L$^{+}$) splenocytes from *Tyk2*$^{+/-}$ (n=11) or *Tyk2*$^{-/-}$ (n=10) 14w mice gated on CD4 or CD8 T cells. (E) Expression heatmap of the whole set of genes in purified β-cells. The arrows indicate the *Insulin I* and *Insulin II* genes, confirming the characteristics of β-cells. (F) Gene ontology (GO) biological process analysis of β-cells from 6w mice.

**Figure S2. Immune cell profiles in the thymus, spleen, PLN, and pancreas.**
(A) Frequencies of the indicated immune cells in the thymi of 6w (*Tyk2*$^{+/-}$, n=9; *Tyk2*$^{-/-}$, n=7), 11w (*Tyk2*$^{+/-}$, n=9; *Tyk2*$^{-/-}$, n=9), and 14w (*Tyk2*$^{+/-}$, n=11; *Tyk2*$^{-/-}$, n=11) NOD mice. (B) Cell number (#) or frequency among CD45$^{+}$ cells (%) of the indicated immune cells in the spleens of 6w (*Tyk2*$^{+/-}$, n=9; *Tyk2*$^{-/-}$, n=7), 11w (*Tyk2*$^{+/-}$, n=9; *Tyk2*$^{-/-}$, n=9), and 14w (*Tyk2*$^{+/-}$, n=12; *Tyk2*$^{-/-}$, n=10) NOD mice. (C) Gating strategy for the analysis of immune cells in the pancreas. The method described in the Materials and Methods section preserves the epitope integrity in immune cells derived from the pancreas. (D, E, F) Cell

number (#) or frequency among CD45$^+$ cells (%) of CD4 T cells (D), γδ T cells (E), and B cells (F) in the PLN of 6w (*Tyk2*$^{+/-}$, n=9; *Tyk2*$^{-/-}$, n=7), 11w (*Tyk2*$^{+/-}$, n=11; *Tyk2*$^{-/-}$, n=11), 14w (*Tyk2*$^{+/-}$, n=11; *Tyk2*$^{-/-}$, n=11), and 24w (*Tyk2*$^{-/-}$, n=3) NOD mice and pancreas of 6w (*Tyk2*$^{+/-}$, n=6; *Tyk2*$^{-/-}$, n=6), 11w (*Tyk2*$^{+/-}$, n=5; *Tyk2*$^{-/-}$, n=9), 14w (*Tyk2*$^{+/-}$, n=11; *Tyk2*$^{-/-}$, n=15), and 24w (*Tyk2*$^{-/-}$, n=8) NOD mice. *P*-values were calculated using unpaired *t*-tests.

**Figure S3. Characteristics of CD4 and CD8 T cells in the PLN and pancreas.**

(A) Frequency of naïve (CD44$^{lo}$ CD62L$^+$), effector memory (CD44$^{hi}$ CD62L$^-$), and central memory (CD44$^{hi}$ CD62L$^+$) CD4 T cells in the pancreas in 6w (*Tyk2*$^{+/-}$, n=4; *Tyk2*$^{-/-}$, n=4), 11w (*Tyk2*$^{+/-}$, n=5; *Tyk2*$^{-/-}$, n=7), 14w (*Tyk2*$^{+/-}$, n=8; *Tyk2*$^{-/-}$, n=12) NOD mice, and CD8 T cells in the pancreas of 6w (*Tyk2*$^{+/-}$, n=5; *Tyk2*$^{-/-}$, n=6), 11w (*Tyk2*$^{+/-}$, n=5; *Tyk2*$^{-/-}$, n=7), and 14w (*Tyk2*$^{+/-}$, n=8; *Tyk2*$^{-/-}$, n=12) NOD mice. (B) Frequency of IFN-γ or IL-17A-producing cells in the pancreas of 11w (*Tyk2*$^{+/-}$, n=5; *Tyk2*$^{-/-}$, n=4 or 5) and 14w NOD mice (*Tyk2*$^{+/-}$, n=8 or 11; *Tyk2*$^{-/-}$, n=12 or 15). (C) Frequency of naïve (CD44$^{lo}$ CD62L$^+$), effector memory (CD44$^{hi}$ CD62L$^-$) and central memory (CD44$^{hi}$ CD62L$^+$) CD4 T cells in the PLN of 6w (*Tyk2*$^{+/-}$, n=6; *Tyk2*$^{-/-}$, n=6), 11w (*Tyk2*$^{+/-}$, n=6; *Tyk2*$^{-/-}$, n=7), and 14w (*Tyk2*$^{+/-}$, n=18; *Tyk2*$^{-/-}$, n=15) NOD mice, and CD8 T cells in the PLN of 11w (*Tyk2*$^{+/-}$, n=6; *Tyk2*$^{-/-}$, n=7) and 14w (*Tyk2*$^{+/-}$, n=18; *Tyk2*$^{-/-}$, n=15) NOD mice. (D) Frequency of naïve (CD44$^{lo}$ CD62L$^+$), effector memory (CD44$^{hi}$ CD62L$^-$), and central memory (CD44$^{hi}$ CD62L$^+$) CD4 T cells in the iLN of 6w NOD mice (*Tyk2*$^{+/-}$, n=6; *Tyk2*$^{-/-}$, n=5), and CD8 T cells in the iLN of 6w NOD mice (*Tyk2*$^{+/-}$, n=6; *Tyk2*$^{-/-}$, n=6). (E) Frequency of IFN-γ or IL-17A-producing cells in the iLN of 6w NOD mice (*Tyk2*$^{+/-}$, n=4; *Tyk2*$^{-/-}$, n=5). (F) Flow cytometry plots of IFN-γ and IL-17-producing CD4 T cells in the PLN of 6w NOD mice. The graphs indicate the frequency of IFN-γ and IL-17-producing CD4 T cells among CD4 T cells in the PLN of mice at the indicated ages. (G) Frequency of Tregs in the PLN of 6–7w (*Tyk2*$^{+/-}$, n=6; *Tyk2*$^{-/-}$, n=6) and 10–12w (*Tyk2*$^{+/-}$, n=7; *Tyk2*$^{-/-}$, n=6) NOD mice. *P*-values were calculated using unpaired *t*-tests.

**Figure S4. Characteristics of CD8 T cells and DCs in PLN.**

(A) Flow cytometry plots of *Tyk2*$^{+/-}$ or *Tyk2*$^{-/-}$ CTV-labeled 8.3 CD8 T cells in PLN 5d post transfer. (B) Frequency of proliferating *Tyk2*$^{+/-}$ or *Tyk2*$^{-/-}$ CTV-labeled 8.3 CD8 T cells in the iLN of the indicated recipient mice. (C) Flow cytometry plots of DCs (resident DC, MHC $II^{mid}$ $CD11c^{hi}$; migratory DC, MHC $II^{hi}$ $CD11c^{hi}$) in the PLN of 6w mice. (D) Flow cytometry plots of migratory DC in the PLN of 6w mice. The frequency of $CD103^{+}$ $CD11b^{-}$ mDC, $CD103^{+}$ $CD11b^{+}$ mDC, and $CD103^{-}$ $CD11b^{+}$ mDC among mDCs is shown (*Tyk2*$^{+/+}$, n=4; *Tyk2*$^{+/-}$, n=6; *Tyk2*$^{-/-}$, n=4). (E) Expression levels of MHC II (I-$A^{g7}$) in DCs in the PLN of 6w mice (*Tyk2*$^{+/+}$, n=4; *Tyk2*$^{+/-}$, n=6; *Tyk2*$^{-/-}$, n=4). (F) Expression levels of MHC I (H2-$K^{d}$) in DCs in the PLN of 6w mice (*Tyk2*$^{+/+}$, n=4; *Tyk2*$^{+/-}$, n=6; *Tyk2*$^{-/-}$, n=4). (G) Expression levels of MHC I (H2-$K^{d}$) in DCs in the iLN or spleens of 6w mice (*Tyk2*$^{+/+}$, n=5; *Tyk2*$^{+/-}$, n=6; *Tyk2*$^{-/-}$, n=5). (H) Expression levels of CD40 or CD86 in $CD8^{+}$ rDCs in the PLN of 6w mice (*Tyk2*$^{+/+}$, n=5; *Tyk2*$^{+/-}$, n=5; *Tyk2*$^{-/-}$, n=4). (I) Gene ontology (GO) biological process analysis of $Tyk2^{+/+}$ $CD8^{+}$ rDC in PLN. *P*-values were calculated using unpaired *t*-tests. (J) Frequency of $CD44^{hi}$ Eomes (*Tyk2*$^{+/-}$, n=11; *Tyk2*$^{-/-}$, n=9), TCF1 (*Tyk2*$^{+/-}$, n=6; *Tyk2*$^{-/-}$, n=6), or Blimp1 (*Tyk2*$^{+/-}$, n=5; *Tyk2*$^{-/-}$, n=4) -positive CD8 T cells among polyclonal CD8 T cells in the PLN of 6w mice. (K) Frequency of $CD44^{hi}$ Eomes (*Tyk2*$^{+/-}$, n=11; *Tyk2*$^{-/-}$, n=9), TCF1 (*Tyk2*$^{+/-}$, n=6; *Tyk2*$^{-/-}$, n=6), or Blimp1 (*Tyk2*$^{+/-}$, n=5; *Tyk2*$^{-/-}$, n=4) -positive CD8 T cells among IGRP-specific CD8 T cells in the PLN of 6w mice. (L) Frequency of polyclonal $CD44^{hi}$ $Cxcr3^{+}$ CD4 T cells among CD4 T cells or polyclonal $CD44^{hi}$ $Cxcr3^{+}$ CD8 T cells among CD8 T cells in the PLN (CD4 T cells from 6w (*Tyk2*$^{+/+}$, n=7; *Tyk2*$^{+/-}$, n=5; *Tyk2*$^{-/-}$, n=7); 11w (*Tyk2*$^{+/+}$, n=3; *Tyk2*$^{+/-}$, n=4; *Tyk2*$^{-/-}$, n=8); and 14w (*Tyk2*$^{+/+}$, n=3; *Tyk2*$^{+/-}$, n=4; *Tyk2*$^{-/-}$, n=7) mice; CD8 T cells from 6w (*Tyk2*$^{+/+}$, n=7; *Tyk2*$^{+/-}$, n=5; *Tyk2*$^{-/-}$, n=7); 11w (*Tyk2*$^{+/+}$, n=3; *Tyk2*$^{+/-}$, n=4; *Tyk2*$^{-/-}$, n=8); and 14w (*Tyk2*$^{+/+}$, n=3; *Tyk2*$^{+/-}$, n=7; *Tyk2*$^{-/-}$, n=4) mice). (M) Frequency of polyclonal $CD44^{hi}$ $Cxcr3^{+}$ CD8 T cells among CD8 T cells in the iLN (*Tyk2*$^{+/+}$, n=7; *Tyk2*$^{+/-}$, n=10; *Tyk2*$^{-/-}$, n=7) or spleens of 6w mice (*Tyk2*$^{+/+}$, n=5; *Tyk2*$^{+/-}$, n=8; *Tyk2*$^{-/-}$, n=6).

**Figure S5. Effects of BMS-986165 treatment on CTLs, CD4 T cells, and NIT-1 cells.** (A) Expression levels of T-bet and Cxcr3 in *ex vivo*-stimulated CD8 T cells treated with BMS-986165 or vehicle for 2 days after stimulation. (B, C) Frequency of $CD44^{hi}$ IFN-$\gamma^{+}$ cells among CD8 T cells (B) and CD4 T cells (C) induced by IL-12 and IL-18 in the presence of BMS-986165. (D, E) Expression levels of *Cxcl9* (D) and *Isg15* (E) in NIT-1 cells stimulated with IFN-β or INF-γ in the presence or absence of BMS-986165 for 3 hours. *P*-values were calculated using unpaired *t*-tests. The data represent two independent experiments with similar results. The data represent two independent experiments with similar results.

**SUPPLEMENTARY FIGURE**

Figure S1

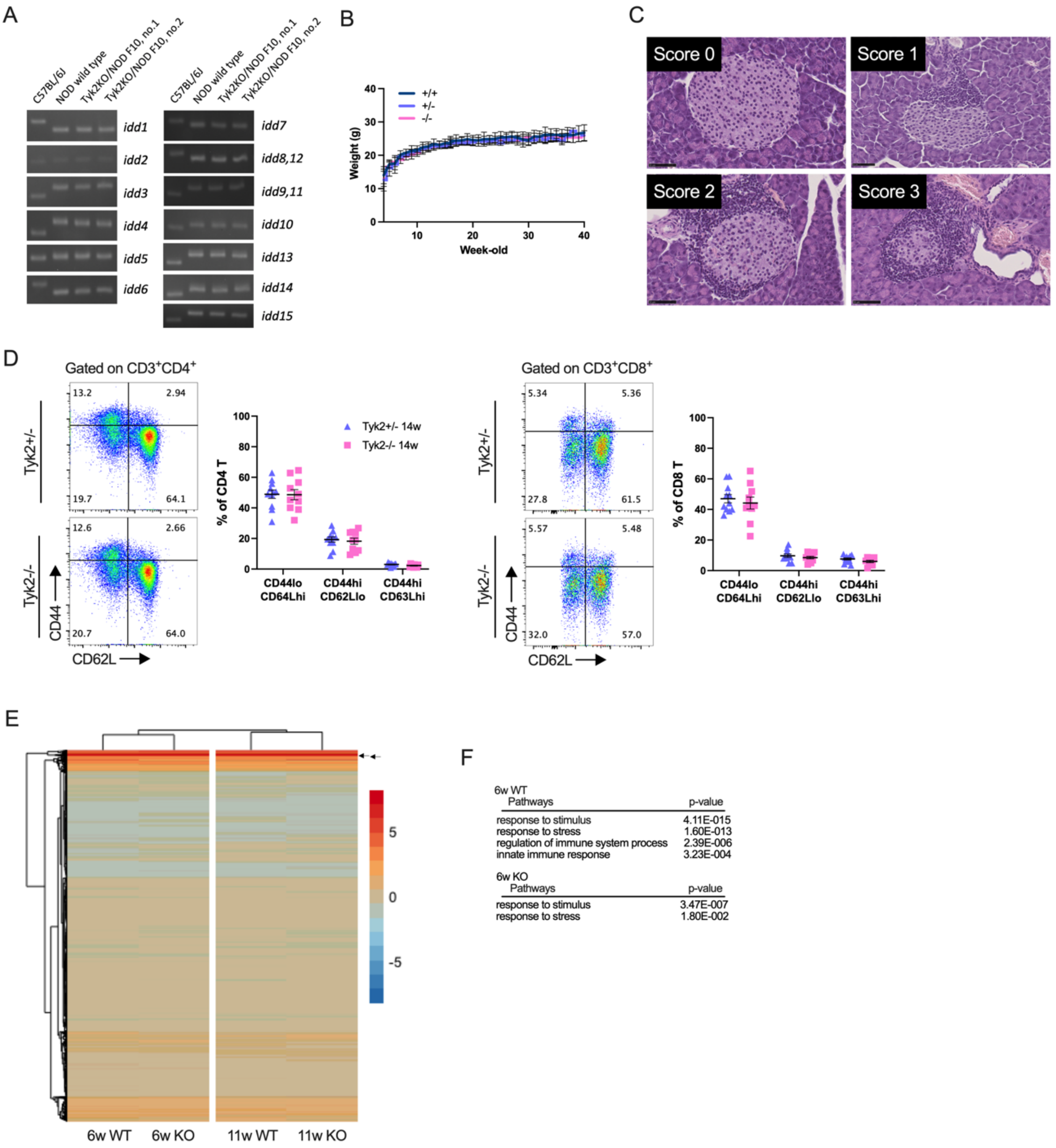


Figure S1. Establishing *Tyk2*KO/NOD mice and the analysis of purified β-cells. Related to Figure 1 and 2.

Figure S2

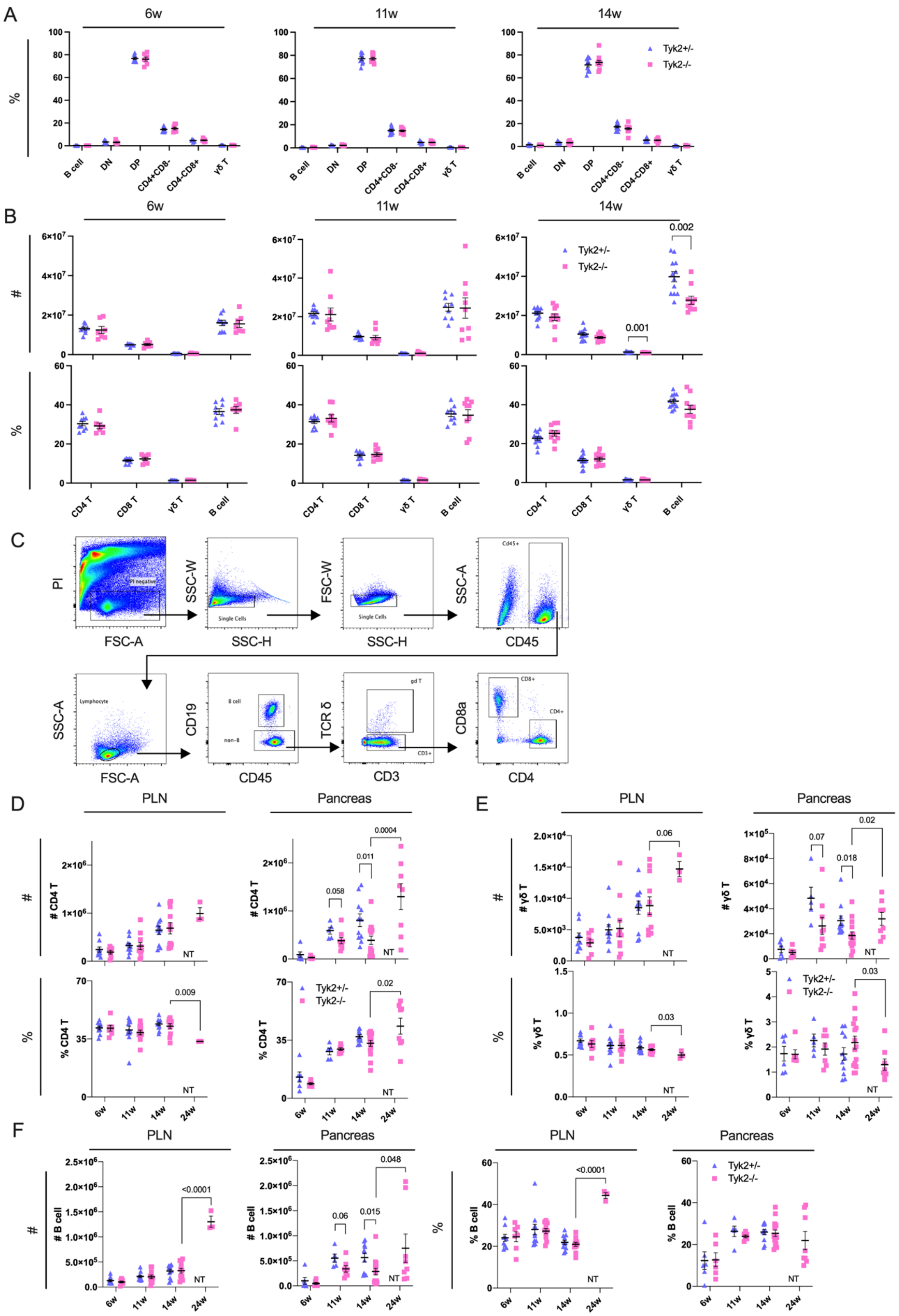

Figure S2. Immune cell profiles in the thymus, spleen, PLN, and pancreas.
Related to Figure 3.

Figure S3

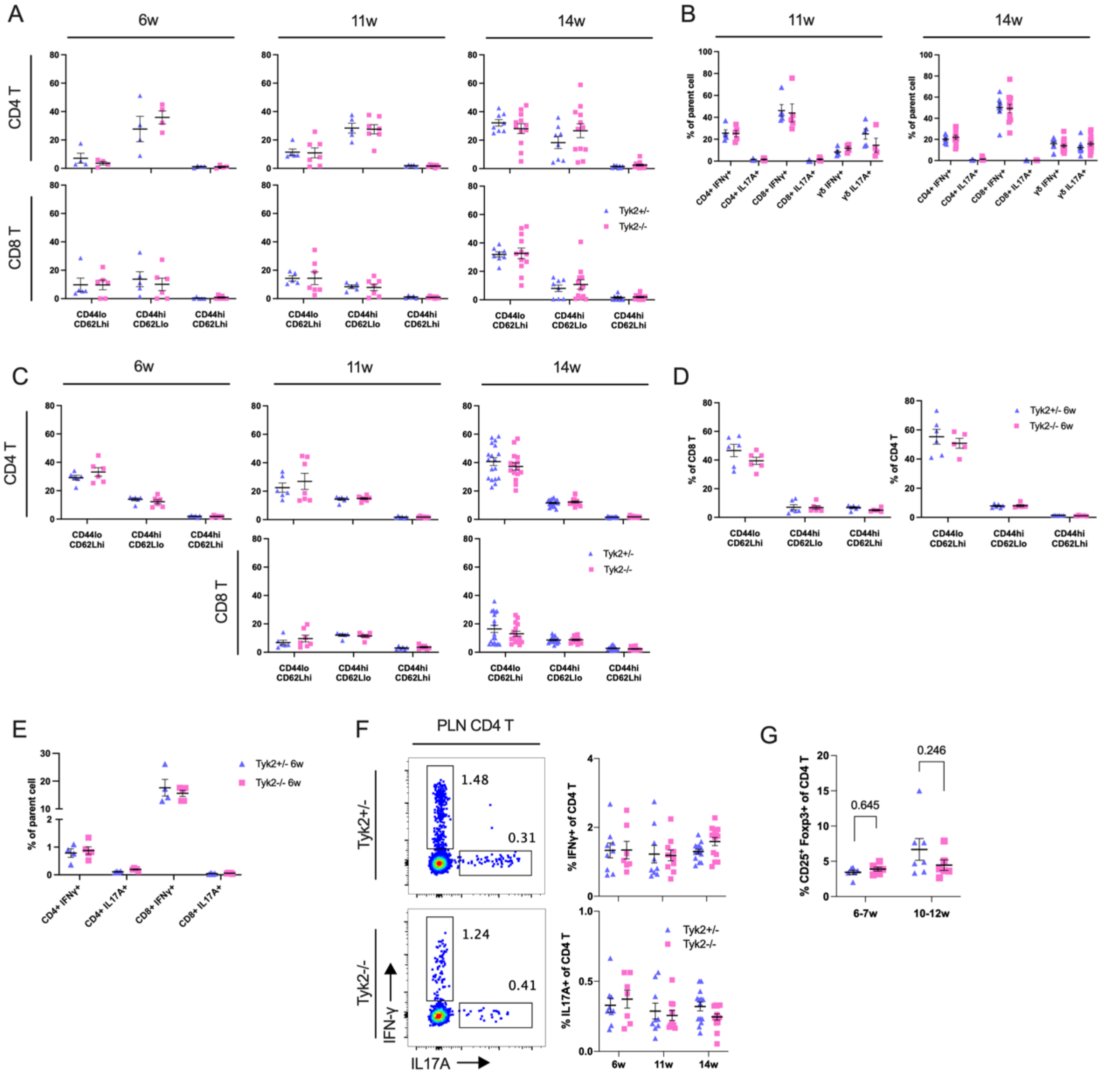


Figure S3. Characteristics of CD4 and CD8 T cells in the PLN and pancreas. Related to Figure 3.

Figure S4

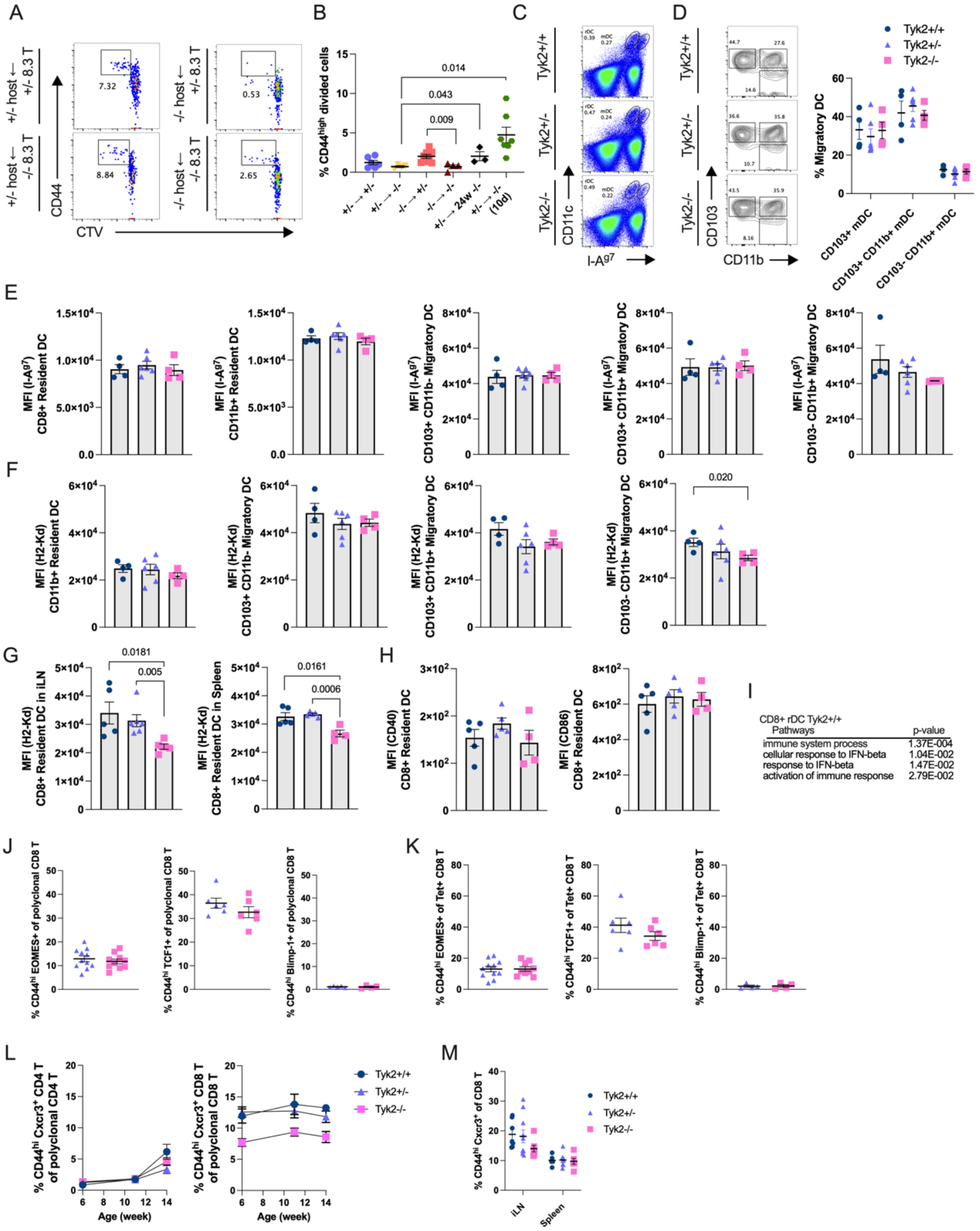


Figure S4. Characteristics of CD8 T cells and DCs in PLN.
Related to Figure 4, 5.

Figure S5

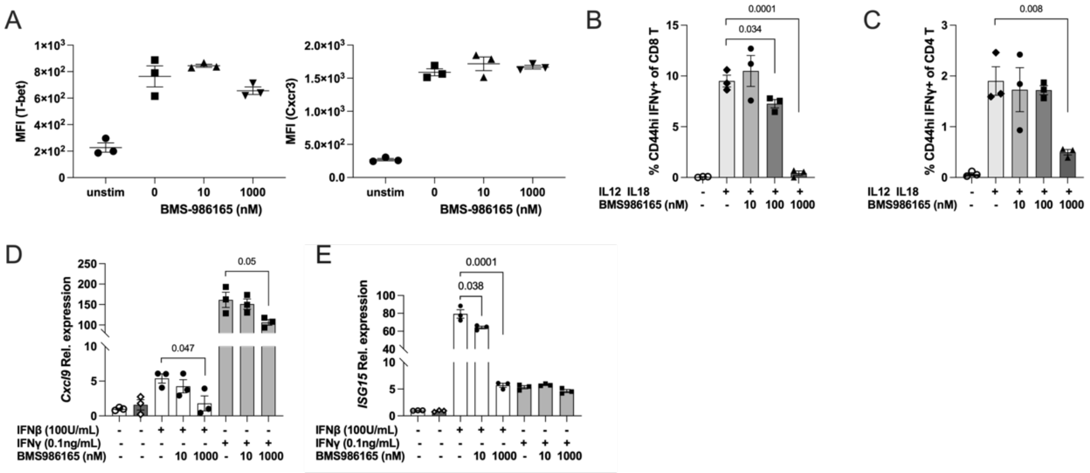


Figure S5. Effects of BMS-986165 treatment on CTLs, CD 4 T cells, and NIT-1 cells. Related to Figure 6.
